# Structural basis for interaction between CLAMP and MSL2 proteins involved in the specific recruitment of the dosage compensation complex in Drosophila

**DOI:** 10.1101/2022.02.16.480628

**Authors:** Evgeniya Tikhonova, Sofia Mariasina, Sergey Efimov, Vladimir Polshakov, Oksana Maksimenko, Pavel Georgiev, Artem Bonchuk

**Author notes:** These authors contributed equally to the paper as first authors (Evgeniya Tikhonova, Sofia Mariasina).

## Abstract

Transcriptional regulators select their targets from a large pool of similar genomic sites. The binding of the *Drosophila* dosage compensation complex (DCC) exclusively to the male X chromosome provides insight into binding site selectivity rules. Previous studies showed that the male-specific organizer of the complex, MSL2, and ubiquitous DNA-binding protein CLAMP directly interact and play an important role in the specificity of X chromosome binding. Here we studied the highly specific interaction between the intrinsically disordered region of MSL2 and the N-terminal zinc-finger C2H2-type (C2H2) domain of CLAMP. We obtained the NMR structure of the CLAMP N-terminal C2H2 zinc finger, which has a classic C2H2 zinc-finger fold with a rather unusual distribution of residues typically used in DNA recognition. Substitutions of residues in this C2H2 domain had the same effect on the viability of males and females, suggesting that it plays a general role in CLAMP activity. The N-terminal C2H2 domain of CLAMP is highly conserved in insects. However, the MSL2 region involved in the interaction is conserved only within the *Drosophila* genus, suggesting that this interaction emerged during the evolution of a mechanism for the specific recruitment of the DCC on the male X chromosome in Drosophilidae.

## INTRODUCTION

It remains unknown how transcription complexes bind exclusively to the certain chromatin regions that do not have pronounced sequence specificity relative to many other genome regions. A striking example of the specific recruitment of transcription complexes is the process of dosage compensation in *Drosophila* (1-3). Dosage compensation occurs by increasing the level of gene expression of the X chromosome of males (X / Y) relative to that of females (X / X). The dosage compensation complex (DCC), which binds only to the male X-chromosome, is responsible for increasing the expression of male genes.

The DCC consists of five proteins, MSL1, MSL2, MSL3, MOF, and MLE, and includes two non-coding RNAs, roX1 (3.7 kb) and roX2 (0.6 kb), which perform mutually interchangeable functions (1,2). Proteins MSL1, MSL3, MOF, and MLE are also present in females and are involved in regulating gene expression in other transcriptional complexes unrelated to dose compensation (1). The MSL2 protein is specific for males (4), and is therefore believed to play a major role in the selective recognition of the male X chromosome.

Inactivation of the MSL3 or MLE proteins led to binding of the incomplete MSL1-MSL2 complex to approximately 200 sites on the X chromosome, referred to as chromatin entry sites (CES) or high-affinity sites (HAS) (5,6). The CXC domain of MSL2 recognizes most HAS/CES sites *in vitro*, and it is likely to be involved in the selection of the male X chromosome by DCC (7-9) In addition, the zinc-finger CLAMP protein binds to GA-repeats found in most of HAS/CES and is essential for recruitment of DCC on the male X chromosome (10). The N-terminal 153 aa region of CLAMP, including the first zinc-finger C2H2 domain, interacts with the unstructured highly conserved (within the *Drosophila* genus) 618−655 aa region of MSL2 (the CLAMP-binding-domain, CBD) (11,12). Inactivation of CBD or CXC in MSL2 only modestly affects recruitment of the DCC to the X chromosome in males (12). However, combining of these two genetic lesions within the same MSL2 mutant resulted in strong inactivation of DCC.

Our previous data suggest that the C2H2 zinc finger domain is required for this interaction (12). The typical C2H2 zinc finger domain has two beta strands and an alpha-helix stabilized by a coordinated zinc ion. Domains of this type are commonly involved in specific DNA binding through residues in the alpha-helix. However, there is growing evidence of their possible function as protein-protein interaction domains (13).

Interaction of the CLAMP N-terminal zinc-finger with MSL2 is highly specific: in Y2H screening, we revealed that MSL2 only interacted with CLAMP and not with any other protein of the tested 152 multi-zinc-finger *Drosophila* transcription factors (14). Here we examined the structural basis for the interaction between the CLAMP and MSL2 proteins using NMR techniques and mutagenic screening complemented with *in vivo* experiments. This study reveals for the first time the features of stable C2H2 zinc-finger interaction with unfolded peptides. We found that the CLAMP N-terminal C2H2 domain interacts with MSL2 using amino acid residues different from those commonly used for DNA recognition. Furthermore, only simultaneous substitutions of several residues at the binding interface significantly weakened the interaction and resulted in progressive DCC delocalization.

## MATERIALS AND METHODS

### Plasmids and cloning

cDNAs were PCR-amplified using corresponding primers (Supplementary Table S3) and cloned into a modified pGEX4T1 vector (GE Healthcare) encoding the TEV protease cleavage site after GST and into the vector derived from pACYC and pET28a(+) (Novagen) bearing a p15A replication origin, kanamycin resistance gene, and pET28a(+) MCS. *Apis mellifera* cDNA was prepared using standard procedures from adult bees obtained from a local apiary. PCR-directed mutagenesis was used to create constructs expressing mutant proteins using mutagenic primers (Supplementary Table S3). For yeast two-hybrid assays, cDNAs were amplified using the corresponding primers (Supplementary Table S3) and fused with the DNA-binding or activation domain of GAL4 in the corresponding pGBT9 and pGAD424 vectors (Clontech). Details of assembling the constructs for expressing proteins in transgenic flies are available upon request.

### Analysis of C2H2 zinc-finger amino acid composition

The development of a hidden Markov Model of the C2H2 domain sequence and calculation of the probabilities of each residue at given positions were performed using Skylign (15). Details are described in Supplementary Information.

### Protein procedures

Proteins were expressed in BL21 (DE3) cells and purified with IMAC, followed by anion-exchange chromatography. Pulldown assays were performed as described (12). Stable isotope-labeled proteins were expressed according to (16) and purified using the same procedures as native proteins. Detailed procedures are described in Supplementary Information.

### NMR spectroscopy

The NMR samples were prepared in concentrations of 0.5 mM (^13^C, ^15^N-labeled MSL2^618-655^ and CLAMP derivatives) and 0.1–0.4 mM (^15^N-labeled protein) with 5% (v/v) D_2_O for frequency lock. Most 2D NMR spectra were collected using a Bruker AVANCE 600 MHz spectrometer equipped with TXI triple resonance (^1^H,^13^N,^15^N) probe, and 3D spectra were recorded on a Bruker AVANCE 700 MHz spectrometer equipped with a quadruple resonance (^1^H, ^13^C, ^15^N, ^31^P) cryo-probe. All NMR experiments were carried out at 25°C. Additional experimental details are included in Supplementary. The structure of CLAMP was deposited to protein data bank with the accession ID 7NF9.

### Y2H

The yeast two-hybrid assay was performed as previously described (17). Briefly, for growth assays, plasmids were transformed into yeast strain pJ69-4A by the lithium acetate method, following standard Clontech protocol, and plated on media without tryptophan and leucine. After two days of growth at 30°C, the cells were plated on selective media without tryptophan, leucine, histidine, and adenine, and their growth was compared after 2–3 days. Each assay was repeated three times.

### Fly crosses, transgenic lines, and polytene chromosome immunostaining

Fly protein extracts were obtained as described (18). Immunostaining of polytene chromosomes was performed as described (12). For details see Supplementary Information.

## RESULTS

### The MSL2 contact surface of the CLAMP protein

The main goal of the study was to understand how the C2H2 domain of CLAMP specifically interacts with MSL2. The interaction domains have been previously mapped to the 618−655 aa of MSL2 and 87−153 aa of CLAMP (12). Since attempts to obtain a crystal of MSL2/CLAMP complex were unsuccessful, we used NMR techniques to study this complex. The backbone and side-chain resonance assignments have been made for CLAMP^87-153^ using a set of 3D NMR spectra obtained for ^15^N and ^13^C-labelled protein samples. According to observed chemical shifts, the order parameter S^2^ values are high for the residues 126−150, indicating that CLAMP^87-153^ is well structured in this region (Supplementary Figure S1).

A family of 20 NMR structures of CLAMP^87-153^ was calculated (Figure 1A) using a set of 357 distance, 40 dihedral angle, and 2 hydrogen bond restraints (see Supplementary Table S1 for details) and restrained molecular dynamics (MD) protocol as described in Supplementary Information. Solution structure reveals a well-defined zinc-finger domain of C2H2 type on the C-terminus of CLAMP^87-153^ (from F127 to E153), whereas the N-terminal portion (from N87 to S126) is disordered (Figure 1A).

**Figure 1.**
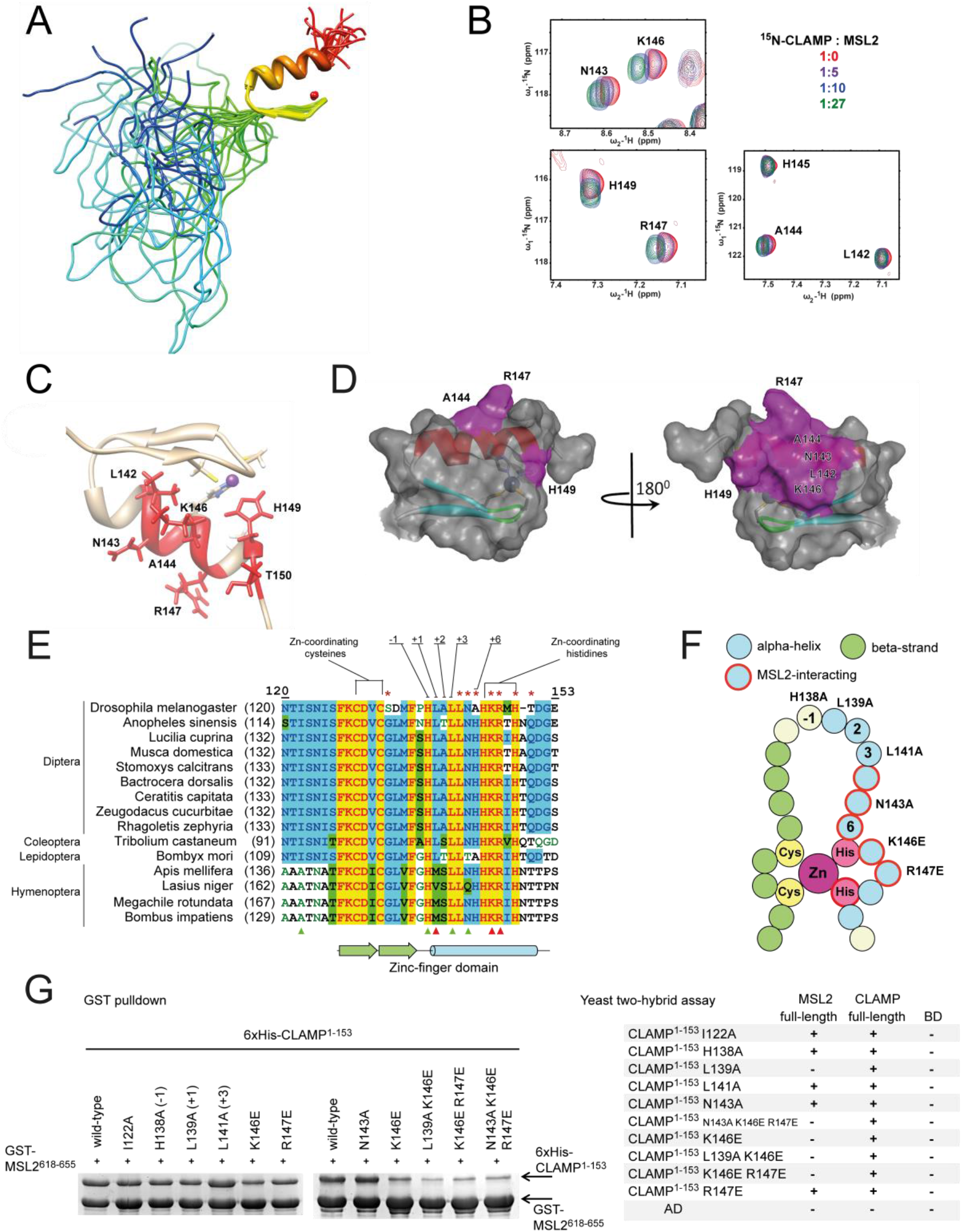
*Drosophila* protein CLAMP interacts with MSL2 through its N-terminal zinc-finger domain. **(A)** NMR structure of CLAMP^87-153^ (PDB 7NF9). **(B)** Sections of ^1^H-^15^N CLAMP^87-153^ spectra showing perturbations of chemical shifts of CLAMP residues upon MSL2 binding. Full spectra are shown in the Supplementary Figure S6. **(C)** Residues with the strongest chemical shifts are shown in red at the CLAMP N-terminal zinc finger structure. **(D)** The MSL2-contact surface of the CLAMP N-terminal zinc finger. **(E)** Multiple sequence alignment of CLAMP N-terminal zinc-fingers from various insects. Typical DNA-binding residues are shown, and red asterisks mark the residues displaying the largest chemical shift perturbations. Triangles depict residues subjected to mutagenesis, red represents a negative effect on binding, and green represents no detectable effect. **(F)** Schematic representation of zinc-finger structure showing the secondary structure, DNA- and MSL2-binding residue positions are depicted. Residues subjected to mutagenesis are shown in red circles. **(G)** GST-pulldown (left) and yeast two-hybrid assays (right) of the interaction between GST-tagged MSL2^618-655^ and 6xHis-thioredoxin-tagged CLAMP^1-153^ bearing point mutations within the zinc-finger domain. AD, activation domain; BD, DNA-binding domain of GAL4 protein. + or - denotes the ability of yeasts to grow on the media without histidine; assay plates are shown in the Supplementary Figure S9.

The CLAMP C2H2 domain structure details are described in Supplementary Figures S2 and S3. The region preceding the zinc finger is conserved in most insects except Hymenopterans (Supplementary Figure S4). CLAMP^1-153^ was found to be able to interact with full-length CLAMP in a yeast two-hybrid assay (Figure 1G) suggesting the presence of oligomerization domain. However, the CLAMP^40-153^ itself does not have dimerization activity in vitro as determined by NMR relaxation studies (Supplementary Figure S5). The superposition of the spectra of CLAMP^87-153^ and CLAMP^1-153^ indicates that the signals of the 126−153 region have the same positions regardless of the lengths of the upstream peptide sequence, while all signals of the 1−125 region are present in the spectral area corresponding to the non-structured polypeptide chain.

To identify critical MSL2-binding residues within CLAMP, we performed NMR chemical shift perturbation experiments with ^15^N-labeled CLAMP^40-153^ titrated with an excess of unlabeled MSL2^618-655^ (Figure 1B and 1C, Supplementary Figure S6). This experiment showed that only the residues within the C2H2 zinc-finger domain exhibit the perturbation of the chemical shifts (Figure 1C and 1D). The most significant changes were found for the residues located within the alpha-helix of the C2H2 domain: N143 (+5 relative to the alpha-helix), T150 (+12), positively charged residues K146 (+8) and R147 (+9), and less for L142 (+4) and A144 (+6). Most of these residues are highly conserved in insects (Figure 1E, Supplementary Figure S4). We also observed chemical shift changes for H149, involved in zinc-ion binding. In this case, the chemical shift perturbation may reflect the slight conformation change of this residue upon binding the MSL2 peptide sequence. Interestingly, most amino acids involved in MSL2 binding differ from those commonly involved in DNA binding by C2H2 zinc fingers (relative to the alpha-helix: -1, +2, +3, +6 and in some cases +1 (19,20)). We compared amino acid residues of the CLAMP N-terminal zinc finger at DNA-binding positions with their average abundance in C2H2 zinc fingers (Supplementary Figure S7). Results of the analysis suggest that the CLAMP N-terminal zinc-finger has quite an atypical pattern of DNA-binding residues. It is very likely, that this zinc-finger is not involved in DNA binding (details provided in Supplementary Information).

To confirm the role of the identified amino acids in MSL2 binding, we tested mutant variants of the C2H2 domain for interaction with MSL2^618-655^ in the GST-pulldown and full-length MSL2 in yeast two-hybrid (Y2H) assays; the ability of CLAMP^1-153^ to interact with full-length CLAMP served as a control (Figure 1G and Supplementary Figure S9). We engineered single amino acid substitutions of K146 and R147 to oppositely charged glutamate to induce electrostatic repulsion and N143 to alanine. Also, we mutated conserved residues at DNA-binding positions H138 (−1), L139 (+1), and L141 (+3) to alanines (Figure 1F). Residues +2 and +6 are not conserved in CLAMP proteins (Figure 1E).

Among seven single amino acid substitutions tested, only L139A and K146E considerably affected the interaction of CLAMP^1-153^ and MSL2 in Y2H (Figure 1G). In GST-pulldown, only double and triple mutations in CLAMP^1-153^ displayed strong effects on the interaction with MSL2^618-655^ (Figure 1G). K146E attenuated the in vitro interaction and adding R147E enhanced this effect. However, adding N143A did not have an additive weakening effect on the interaction in combination with other mutations (Figure 1G).

The N-terminal zinc finger of CLAMP is preceded by sequence NTISNIS conserved in most insects, which may contribute to protein-protein binding; however, no changes in chemical shifts were observed for these residues. We introduced an alanine substitution for conservative isoleucine 122 (I122A), but this mutation did not affect the interaction efficiency (Figure 1G).

Altogether, our results suggest that residues in the alpha-helix of CLAMP N-terminal C2H2 domain display redundancy in MSL2 interactions. Only the substitution of several residues involved in the interaction considerably affects the MSL2 binding.

### The CLAMP contact area of the MSL2 protein

We performed a reverse experiment to understand further the mechanism of CLAMP N-terminal C2H2 domain binding to the unfolded MSL2 peptide. We produced stable isotope-labeled MSL2^618-655^ protein and performed a backbone resonance assignment using a set of 3D NMR spectra on ^15^N and ^13^C-labelled protein samples. Molecular modeling suggests a possibility to form β-hairpins at V634−N638 and G641−N647, but according to the chemical shift values, MSL2^618-655^ is mostly unstructured in solution (Supplementary Figure S10), which is also supported by the lack of NOE signals. Upon titration with unlabeled CLAMP^40-153^ we observed chemical shift perturbation of the certain residues of MSL2^618-655^ (Figure 2A, relative perturbation is shown in Figure 2B, full spectra in Supplementary Figure S11). We did not observe a significant change in chemical shifts of MSL2^618-655^ upon binding to CLAMP, which indicates that it does not acquire ordered conformation in the process of complex formation.

**Figure 2.**
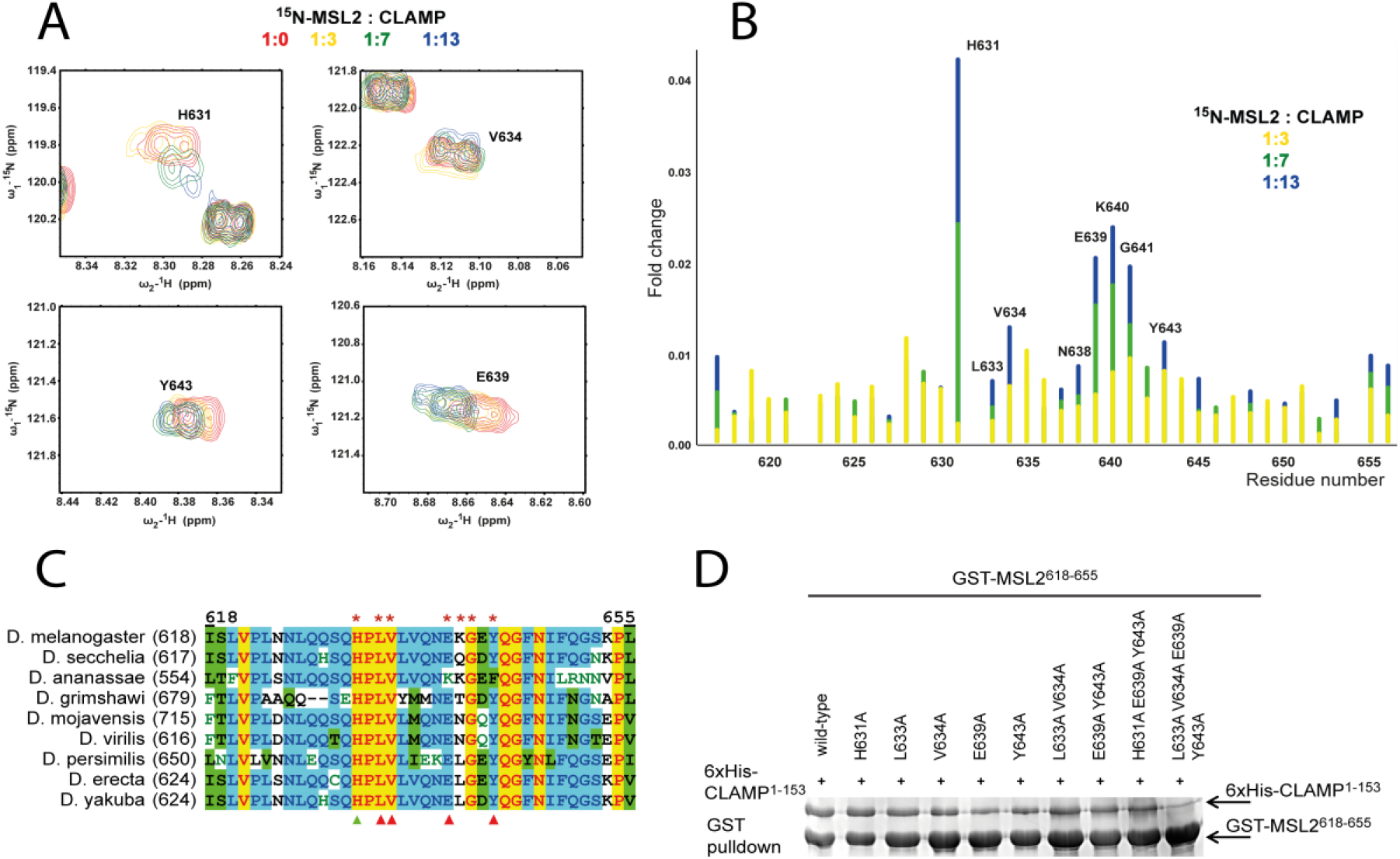
The small unfolded sequence of MSL2 interacts with CLAMP. **(A)** Sections of ^1^H-^15^N MSL2^618-655^ spectra showing perturbations of chemical shifts of MSL2 residues upon CLAMP binding. Full spectra are shown in the Supplementary Figure S11. **(B)** Relative perturbation of MSL2 residues’ chemical shifts upon interaction with CLAMP. **(C)** Multiple sequence alignment of CLAMP-interacting MSL2 region from *Drosophila* species. Red asterisks show the residues with the largest chemical shift perturbations upon binding to CLAMP. Triangles mark residues subjected for mutagenesis, red represents a negative effect on binding, and green represents no detectable effect. **(D)** GST-pulldown assay of interaction between GST-tagged MSL2^618-655^ (represented by *) and 6xHis-thioredoxin-tagged CLAMP^1-153^.

The NMR spectra measured during titration experiments indicate a fast exchange between free and bound states, which is a consequence of relatively weak protein-protein interaction. For fast exchange between bound and free states (k_off_ > R_1_), resonances of labeled protein moved from free to bound positions upon the increase of concentration of unlabeled interacting protein. Analysis of chemical shift perturbation during the titration allowed to estimate the K_d_ of CLAMP-MSL2 interaction as 0.22 ± 0.13 mM at 25°C (see Supplementary Information for details).

Largest chemical shift changes were observed for both hydrophobic (L633, V634, Y643) and charged clusters of residues (E639, K640), with the most significant change for H631. To further demonstrate the influence of these residues on CLAMP binding, we designed substitutions for conserved residues H631A, L633A, V634A, E639A, and Y643A. We utilized combinations of these substitutions and introduced them simultaneously. Unexpectedly, H631A did not affect the interaction even though it displayed the strongest chemical shift perturbation. L633A and V634A had little effect, but E639A, Y643A, and their combination significantly weakened binding of MSL2 to CLAMP *in vitro*. Furthermore, simultaneous substitutions of four residues (L633A, V634A, E639A and Y643A) resulted in almost complete loss of interaction in the pulldown assay (Figure 2D). These results suggest that the interaction between MSL2 and CLAMP is cooperative and allows in the course of evolution to gradually select the most effective combination of amino acids in MSL2 for interaction with the CLAMP N-terminal C2H2 domain.

Almost all residues responsible for the interaction with CLAMP are highly conserved in the MSL2 proteins of various Drosophilids (Figure 2C). However, the full motif is not conserved even in the closest Dipterans (Supplementary Figure S12). This low conservation contrasts with the CLAMP C2H2 domain, which remains highly conserved in all insects. Bees and other social Hymenopterans have a completely different mechanism of sex determination (males are haploid) and thus might not utilize dosage compensation similar to Drosophilids (21). At the same time, the N-terminal C2H2 domain has a high level of homology between CLAMP proteins from *Drosophila melanogaster* and *Apis mellifera*. We studied CLAMP-MSL2 interaction in *Apis mellifera* (hereafter amCLAMP and amMSL2). The amCLAMP and amMSL2 do not interact directly (Supplementary Figure S13B). At the same time, amCLAMP can interact with *D. melanogaster* MSL2^618-655^, which is not surprising since most protein-interacting residues are conserved (Figure 1E, Supplementary Figure S13).

### Mutations of the MSL2-CLAMP interface affect DCC recruitment *in vivo*

To assess the effect of mutations disrupting the CLAMP:MSL2 contact surface *in vivo*, we generated transgenic flies expressing 3xHA-tagged CLAMP^WT^, CLAMP^K146E^, CLAMP^K146E;R147E^, and CLAMP^L139A;K146E^ under strong ubiquitin (*Ubi63E*) promoter (Figure 3A). The transgenes were inserted into the same 86Fb region on the third chromosome, using a φC31 integrase-based integration system (22). Immunoblot analysis showed that all CLAMP variants were expressed at similar levels (Supplementary Figure S14A). To understand the functional roles of the mutations in the C2H2 domain, we used a previously described null mutation in the *clamp* gene, named *clamp*^*2*^ (23). We examined the ability of transgenes expressing wild-type and mutant proteins to complement the *clamp*^*2*^ mutation (Figure 3B, Supplementary Table S2). Expression of the CLAMP^WT^ protein restored to a greater extent the survival rate of *clamp*^*2*^ flies. At the same time, males and females expressing any of the CLAMP mutants displayed low viability. Thus, the N-terminal domain of C2H2 is required for CLAMP activity, which is not related to its role in dosage compensation.

**Figure 3.**
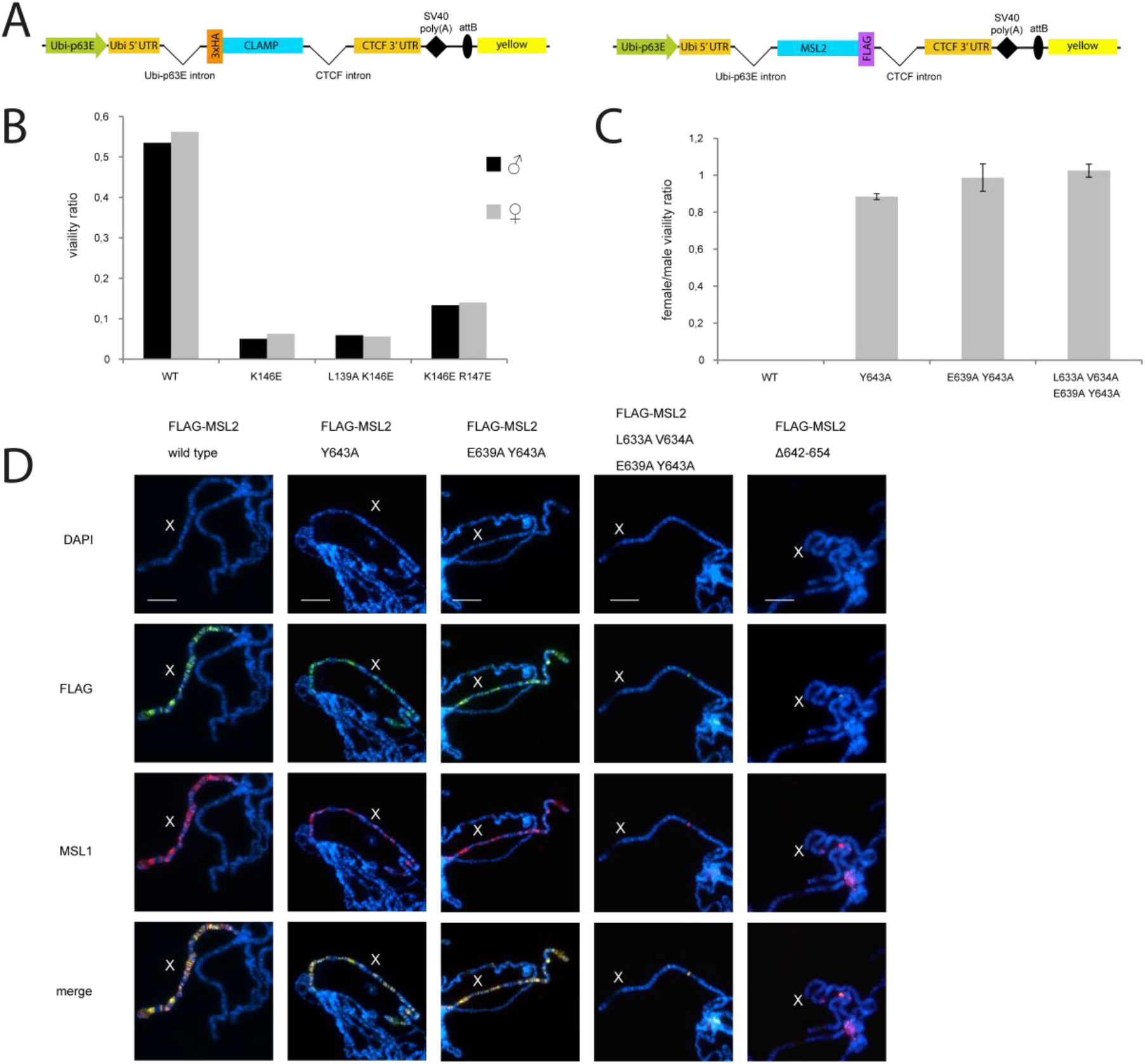
The CLAMP-MSL2 interaction is required for the correct recruitment of the dosage compensation complex. **(A)** Schematic representation of rescue constructs expressing 3xHA-tagged CLAMP 3xFLAG-tagged MSL2 proteins under the control of ubiquitin-p63E promoter; SV40 poly(A)–SV40 polyadenylation signal; attB is the site for φC31-mediated recombination used for site-specific insertion of the construct; yellow represents the intronless *yellow* gene used as a reporter. **(B)** Comparison of the viability (relative to clamp^2^/+ mutant background) of males and females upon the rescue of the clamp^2^/clamp^2^ mutant background with CLAMP proteins expressed in heterozygous (k/TM6) and homozygous (k/k) transgenic constructs. Complete data are shown in the Supplementary Table S2. **(C)** Relative viability of females with ectopic expression of MSL2. **(D)** Effect of single amino-acid substitutions in the FLAG-tagged MSL2 proteins on DCC recruitment shown by immunostaining of polytene chromosomes with anti-FLAG and anti-MSL1 antibodies in females. Polytene chromosomes stained with MSL2 antibodies are shown in Supplementary Figure S14B. Scale bar is 20 µm.

To define the contribution *in vivo* of key amino acids in the CLAMP-interacting region of MSL2, we created transgenic lines expressing different FLAG-tagged MSL2 mutants under control of the *Ubi63E* promoter in the 86Fb region (Figure 3A). We tested the mutations in the MSL2 that affect interaction with CLAMP *in vitro*: MSL2^Y643A^, MSL2^E639A;Y643A^, and MSL2^L633A;V634A;E639A;Y643A^. The previously obtained transgenic lines (12) expressing MSL2^WT^ and MSL2^Δ13d^ (deletion of CLAMP interacting region, 642−654 aa, in MSL2) were used as positive and negative controls. Immunoblot analysis showed that all MSL2 variants were present in the transgenic flies at nearly equivalent levels as MSL2^WT^ (Supplementary Figure S14A).

Ectopic expression of MSL2 in females resulted in the assembly of functional DCC (24,25) that led to a strong increase of gene expression, and as a consequence, decreased viability. This model is most sensitive for identifying mutations in MSL2 that affect the specific binding of the MSL complex to the X chromosome. Females carrying homozygous MSL2^wt^ transgenes had low viability (Figure 3C). At the same time, females carrying homozygous transgenes expressing MSL2^Δ13d^ or any of the tested mutant versions of MSL2 (MSL2^Y643A^, MSL2^E639A;Y643A^, and MSL2^L633A;V634A;E639A;Y643A^) displayed normal viability suggesting that the functional activity of MSL2 is impaired in all mutants (Figure 3C). These results indicate that all mutations in MSL2 negatively influence the assembly of functional DCC in females.

Polytene chromosomes in the nuclei of salivary glands are a well-established model system for studying the recruitment and spreading of the MSL complex along the X chromosome (26-29). We used anti-FLAG mouse antibodies to identify tagged MSL2 variants and anti-MSL1 and anti-MSL2 rabbit antibodies to confirm the recruitment of endogenous components of the DCC (Figure 3D, Supplementary Figure S14B). Transgenic expression of MSL2^WT^ in females led to localization to the X-chromosomes of both the MSL2 and MSL1 proteins (12). The deletions in the CLAMP interacting region resulted in an almost complete absence of binding sites for the MSL2^Δ13d^ mutant and MSL1 on the female X chromosomes. The Y643A and E639A;Y643A mutations in MSL2 only slightly decreased the binding of the mutant MSL2 variants and MSL1 to the female X chromosomes. Simultaneous mutation of four amino acids (L633A;V634A;E639A;Y643A) led to the same effect as the deletion of the CLAMP interacting region.

These results confirm that several residues in MSL2 cooperatively interact with the C2H2 domain of CLAMP. The effectiveness of such interaction depends on the number of residues involved in the formation of a specific CLAMP-MSL2 contact.

## DISCUSSION

For the first time, this study reveals the features of the highly specific interaction of the C2H2 zinc-finger with an intrinsically disordered polypeptide chain – part of MSL2. According to NMR data this fragment of MSL2 does not become structured upon binding to CLAMP. The N-terminal C2H2 domain of CLAMP at DNA-binding positions contains residues that differ significantly from those typical to the C2H2 domain involved in DNA binding. Single amino-acid substitutions have little effect on the CLAMP-MSL2 interaction suggesting an additive role of multiple bonds in forming a stable and specific interaction. Both hydrophilic and hydrophobic residues are involved in the interaction; notably, strong responses were shown for E639 in MSL2 and K146 and R147 in CLAMP, suggesting electrostatic interactions and possible salt bridge formation. Unexpectedly, whereas MSL2 peptide displayed significant chemical shifts for multiple hydrophobic residues after binding to CLAMP, in the CLAMP we can see perturbations mostly for the polar and charged residues. One possible explanation is that the hydrophobic core of the zinc finger did not undergo structural rearrangements after binding the MSL2, but the same residues formed stable hydrophobic pockets involved in the interaction with hydrophobic MSL2 residues.

All residues involved in the interaction are highly conserved in CLAMP proteins but almost non-conserved in family of MSL2 proteins outside the *Drosophila* genus, suggesting that the CLAMP N-terminal zinc finger has an important role and most likely is involved in interactions with other factors besides MSL2. Expression of the mutant CLAMP proteins has an equal effect on male and female viability, further supporting this hypothesis. Even *Apis mellifera* CLAMP can interact with *Drosophila* MSL2, supporting the assumption of strong evolutionary conservation of the CLAMP N-terminal C2H2 domain. However, we did not observe any interaction between amMSL2 and amCLAMP. Thus, it seems likely that the interaction between CLAMP and MSL2 occurs only in Drosophilidae and closely related species since we were unable to identify MSL2 motifs that can interact with the CLAMP C2H2 domain outside the *Drosophila* genus (Supplementary Figure S10).

Previously it was suggested that the interaction between MSL2 and the CLAMP C2H2 domain is essential for recruiting of the DCC to the male X chromosome (12). An alternative proposal was that the roX RNAs bind to the C-terminal portion of MSL2, including the CLAMP interaction region (30,31). The roX RNAs and the MSL2 CTD form a stably condensed state that allows specific recruitment of the MSL complex on the X chromosome. Here we demonstrate that point mutations that impair interaction between CLAMP and MSL2 also affect MSL complex recruitment. It seems unlikely that these point mutations affect roX binding to MSL2. Thus, the CLAMP-MSL2 interaction plays an important role in specific MSL complex recruitment on the X chromosome.

CLAMP is a key early developmental pioneer transcription factor involved in maintaining open chromatin and recruiting major transcription factors (10,23,32,33). During early embryogenesis, CLAMP preferentially binds to genomic regions outside HAS (34). The MSL complex initially binds nonspecifically to CLAMP-rich regions throughout the genome. Subsequently, preferential enrichment of the MSL complex and CLAMP occurs at HAS (11,34). Cooperative interaction with DNA and CLAMP allows MSL to compete more effectively with nucleosomes when binding to chromatin (11). Improving the interaction between MSL2 and CLAMP has an evolutionary advantage as it results in more efficient specific binding of the MSL complex to the male X chromosome. It can be assumed that in the course of evolution, there was a gradual increase in the strength of interaction between CLAMP and MSL2 as a result of mutations in the MSL2 region, which is not conservative.

In *D. virilis*, a species separated from *D. melanogaster* by 40 million years of evolution, orthologs of MSL2 and CLAMP (dvMSL2 and dvCLAMP) interact much more strongly than in *D. melanogaster* (30). However, unlike MSL2, the CXC domain in dvMSL2 does not specifically recognize HAS sites and binds X chromosome and autosomes with the same efficiency. It seems likely that in *D. virilis*, the loss of specific binding of CXC on HAS is compensated by stronger interaction between dvMSL2 and dvCLAMP (Figure 4A). In addition to recruiting the MSL complex, MSL2 binds to autosomal promoters of genes involved in patterning and morphogenesis and is required for the proper development of males (35). It can be assumed that the lack of specificity of the interaction between the CXC domain and the HAS allows dvMSL2 to more efficiently bind to autosomal promoters and participate in their regulation in *D. virilis*.

**Figure 4.**
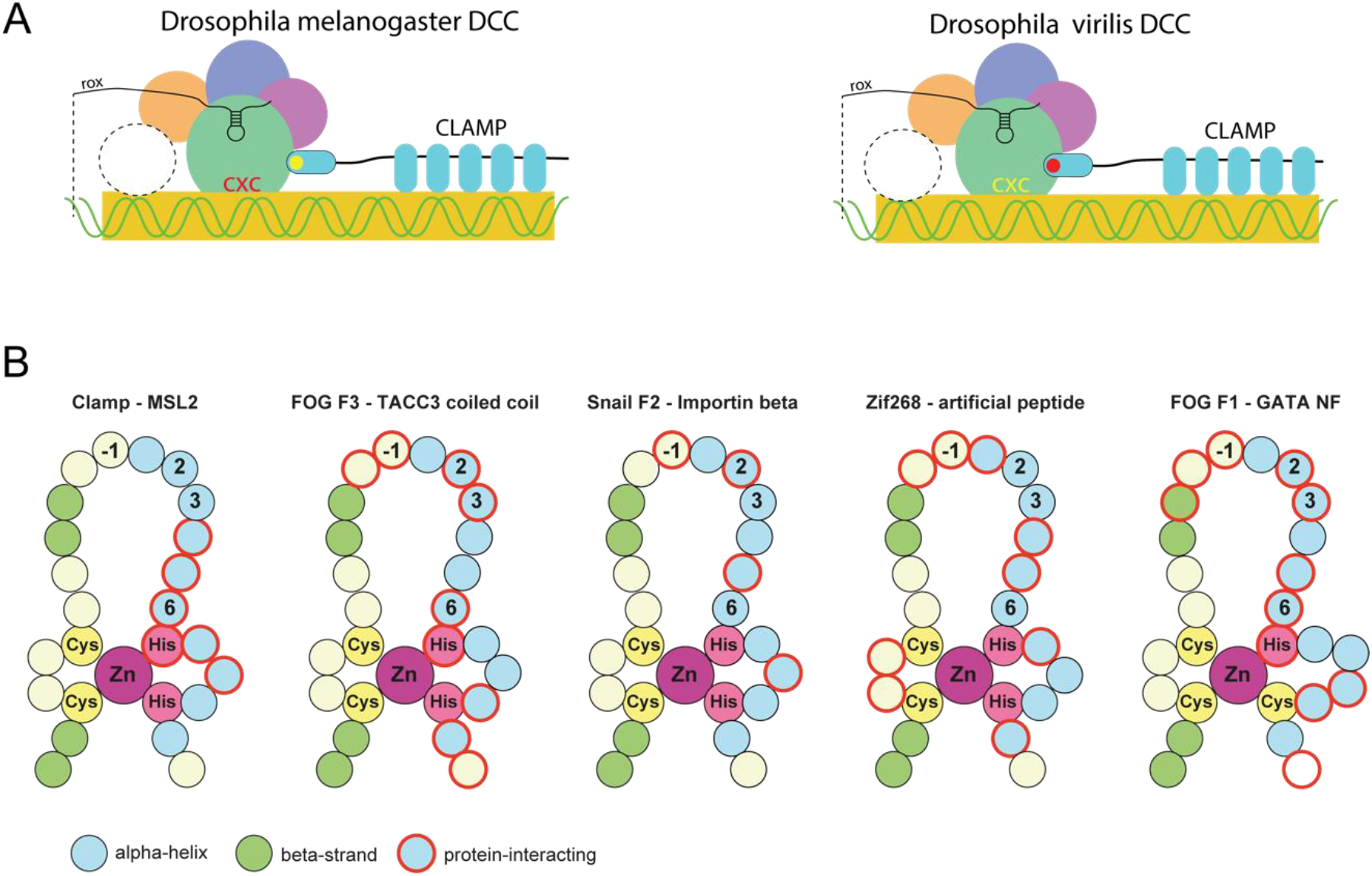
**(A)** Schematic model representing the factor cooperation in the process of specific DCC recruitment to the male X-chromosome. Red color means stronger and yellow indicates weaker interactions. **(B)** Comparison of the zinc-finger residues used for protein binding. Cartoons were drawn according to the following structures (PDB ID): 1F2I (Zif268-artificial peptide), 1Y0J (FOG F1–GATA), 1SRK (FOG F3–TACC3 coiled-coil), 3W5K (Snail F2–importin beta). The cartoons for intramolecular interactions mediated by zinc fingers are shown in the Supplementary Figure S15.

Our results suggest that interactions between C2H2 domains and intrinsically disordered regions may be widespread for creating new protein-protein interactions. However, the availability of structural data supporting the protein binding potential of C2H2 domains is limited. We have summarized the available data for protein-protein interactions mediated by the classical C2H2 domains (Figure 4B and Supplementary Figure S15). The interactions between the C2H2 and CCHC FOG domains with folded protein domains (coiled-coil domain or GATA-type zinc finger) are formed by the alpha-helix, and residues in such interfaces significantly overlap with those involved in DNA-binding (36,37). Zinc fingers of Snail protein also use the DNA-binding side of the alpha-helix for interaction with importin beta (38), but the set of residues significantly differs from that normally used for nucleic acid binding (Figure 4B).

In contrast to these complexes, protein-binding residues of the CLAMP N-terminal C2H2 domain are located closer to the C-terminus and at the opposite side of the alpha-helix. NMR spectra demonstrate that the CLAMP-interacting sequences of MSL2 are unfolded. There is only one example of a similar interaction in which an artificial unfolded peptide contacted with the surface of the alpha-helix opposite to that used for DNA binding in the C2H2 domain of Zif268 (39) (Figure 4B). This interface of C2H2 also was implicated in multiple contacts with structured protein domains. For example, this interface participates in intermolecular interactions between neighbor C2H2 domains of the GLI protein (40,41) and between the first and third C2H2 domains of Kaiso with their additional C-terminal beta-strands (42) (Supplementary Figure S15). A similar interface is used by some closely related UBZ-type C2H2 fingers for ubiquitin recognition, while other domains of this type use only the distal part of alpha-helix (43,44) (Supplementary Figure S15). These examples demonstrate that protein interactions mediated by non-DNA-binding interfaces of classical zinc fingers can be a common property of these domains. Most of the described interactions involve structured protein domains. Therefore, the recognition of an intrinsically disordered MSL2 peptide by a classical C2H2 zinc finger of CLAMP is the only described naturally occurring interaction of this type. Many other classical C2H2 zinc-finger domains can be involved in such interactions.

## ACCESSION NUMBERS

Atomic coordinates and structure factors for the reported NMR structure have been deposited with the Protein Data Bank under accession number 7NF9. The ^1^H, ^15^N and ^13^C chemical shifts of CLAMP^87-153^ have been deposited into the BioMagResBank (www.bmrb.wisc.edu) under the accession number BMRB-34600.

## SUPPLEMENTARY DATA

Supplementary Data are available at NAR online.

## ACKNOWLEDGMENTS

We thank Erica Larschan for the *clamp*^*2*^ mutation, Farhod Hasanov and Aleksander Parshikov for fly injections, and Olga Kyrchanova for assistance with fly genetics. This study was performed using the equipment of the IGB RAS Core Facilities Centre.

## FUNDING

This work was supported in part by the Russian Foundation for Basic Research - grant 19-04-00933 (NMR experiments, genetic and immunostaining experiments), by the Russian Science Foundation - grant 21-14-00211 (expression and purification of proteins and their mutants), Ministry of Science and Higher Education of the Russian Federation — grant 075-15-2019-1661 (analysis of protein-protein interactions) and by Interdisciplinary Scientific and Educational School of Moscow University «Molecular Technologies of the Living Systems and Synthetic Biology» (NMR data analysis).

## CONFLICT OF INTEREST

None declared.

## Notes

### Competing Interest Statement

The authors have declared no competing interest.

## REFERENCES

1. Samata, M. and Akhtar, A. (2018) Dosage Compensation of the X Chromosome: A Complex Epigenetic Assignment Involving Chromatin Regulators and Long Noncoding RNAs. Annu Rev Biochem, 87, 323–350.

2. Kuroda, M.I., Hilfiker, A. and Lucchesi, J.C. (2016) Dosage Compensation in Drosophila-a Model for the Coordinate Regulation of Transcription. Genetics, 204, 435–450.

3. Lucchesi, J.C. (2018) Transcriptional modulation of entire chromosomes: dosage compensation. J Genet, 97, 357–364.

4. Beckmann, K., Grskovic, M., Gebauer, F. and Hentze, M.W. (2005) A dual inhibitory mechanism restricts msl-2 mRNA translation for dosage compensation in Drosophila. Cell, 122, 529–540.

5. Straub, T., Grimaud, C., Gilfillan, G.D., Mitterweger, A. and Becker, P.B. (2008) The chromosomal high-affinity binding sites for the Drosophila dosage compensation complex. PLoS Genet, 4, e1000302.

6. Alekseyenko, A.A., Peng, S., Larschan, E., Gorchakov, A.A., Lee, O.K., Kharchenko, P., McGrath, S.D., Wang, C.I., Mardis, E.R., Park, P.J. et al. (2008) A sequence motif within chromatin entry sites directs MSL establishment on the Drosophila X chromosome. Cell, 134, 599–609.

7. Fauth, T., Muller-Planitz, F., Konig, C., Straub, T. and Becker, P.B. (2010) The DNA binding CXC domain of MSL2 is required for faithful targeting the Dosage Compensation Complex to the X chromosome. Nucleic Acids Res, 38, 3209–3221.

8. Zheng, S., Villa, R., Wang, J., Feng, Y., Becker, P.B. and Ye, K. (2014) Structural basis of X chromosome DNA recognition by the MSL2 CXC domain during Drosophila dosage compensation. Genes Dev, 28, 2652–2662.

9. Villa, R., Schauer, T., Smialowski, P., Straub, T. and Becker, P.B. (2016) PionX sites mark the X chromosome for dosage compensation. Nature, 537, 244–248.

10. Soruco, M.M., Chery, J., Bishop, E.P., Siggers, T., Tolstorukov, M.Y., Leydon, A.R., Sugden, A.U., Goebel, K., Feng, J., Xia, P. et al. (2013) The CLAMP protein links the MSL complex to the X chromosome during Drosophila dosage compensation. Genes Dev, 27, 1551–1556.

11. Albig, C., Tikhonova, E., Krause, S., Maksimenko, O., Regnard, C. and Becker, P.B. (2019) Factor cooperation for chromosome discrimination in Drosophila. Nucleic Acids Res, 47, 1706–1724.

12. Tikhonova, E., Fedotova, A., Bonchuk, A., Mogila, V., Larschan, E.N., Georgiev, P. and Maksimenko, O. (2019) The simultaneous interaction of MSL2 with CLAMP and DNA provides redundancy in the initiation of dosage compensation in Drosophila males. Development, 146.

13. Brayer, K.J., Kulshreshtha, S. and Segal, D.J. (2008) The protein-binding potential of C2H2 zinc finger domains. Cell biochemistry and biophysics, 51, 9–19.

14. Babosha, V., Klimenko, N., Tikhonova, E., Shilovich, A., Georgiev, P. and Maksimenko, O. (2021) Preprint at https://www.biorxiv.org/content/10.1101/2020.11.11.378323v1.

15. Wheeler, T.J., Clements, J. and Finn, R.D. (2014) Skylign: a tool for creating informative, interactive logos representing sequence alignments and profile hidden Markov models. BMC Bioinformatics, 15, 7.

16. Marley, J., Lu, M. and Bracken, C. (2001) A method for efficient isotopic labeling of recombinant proteins. J Biomol NMR, 20, 71–75.

17. Zolotarev, N., Fedotova, A., Kyrchanova, O., Bonchuk, A., Penin, A.A., Lando, A.S., Eliseeva, I.A., Kulakovskiy, I.V., Maksimenko, O. and Georgiev, P. (2016) Architectural proteins Pita, Zw5,and ZIPIC contain homodimerization domain and support specific long-range interactions in Drosophila. Nucleic Acids Res, 44, 7228–7241.

18. Sabirov, M., Kyrchanova, O., Pokholkova, G.V., Bonchuk, A., Klimenko, N., Belova, E., Zhimulev, I.F., Maksimenko, O. and Georgiev, P. (2021) Mechanism and functional role of the interaction between CP190 and the architectural protein Pita in Drosophila melanogaster. Epigenetics Chromatin, 14, 16.

19. Vandevenne, M., Jacques, D.A., Artuz, C., Nguyen, C.D., Kwan, A.H., Segal, D.J., Matthews, J.M., Crossley, M., Guss, J.M. and Mackay, J.P. (2013) New insights into DNA recognition by zinc fingers revealed by structural analysis of the oncoprotein ZNF217. J Biol Chem, 288, 10616–10627.

20. Gupta, A., Christensen, R.G., Bell, H.A., Goodwin, M., Patel, R.Y., Pandey, M., Enuameh, M.S., Rayla, A.L., Zhu, C., Thibodeau-Beganny, S. et al. (2014) An improved predictive recognition model for Cys(2)-His(2) zinc finger proteins. Nucleic Acids Res, 42, 4800–4812.

21. Heimpel, G.E. and de Boer, J.G. (2008) Sex determination in the Hymenoptera. Annu Rev Entomol, 53, 209–230.

22. Bischof, J., Maeda, R.K., Hediger, M., Karch, F. and Basler, K. (2007) An optimized transgenesis system for Drosophila using germ-line-specific phiC31 integrases. Proc Natl Acad Sci U S A, 104, 3312–3317.

23. Urban, J.A., Doherty, C.A., Jordan, W.T., 3rd, Bliss, J.E., Feng, J., Soruco, M.M., Rieder, L.E., Tsiarli, M.A. and Larschan, E.N. (2017) The essential Drosophila CLAMP protein differentially regulates non-coding roX RNAs in male and females. Chromosome Res, 25, 101–113.

24. Kelley, R.L., Wang, J., Bell, L. and Kuroda, M.I. (1997) Sex lethal controls dosage compensation in Drosophila by a non-splicing mechanism. Nature, 387, 195–199.

25. Kelley, R.L., Solovyeva, I., Lyman, L.M., Richman, R., Solovyev, V. and Kuroda, M.I. (1995) Expression of msl-2 causes assembly of dosage compensation regulators on the X chromosomes and female lethality in Drosophila. Cell, 81, 867–877.

26. Zhou, S., Yang, Y., Scott, M.J., Pannuti, A., Fehr, K.C., Eisen, A., Koonin, E.V., Fouts, D.L., Wrightsman, R., Manning, J.E. et al. (1995) Male-specific lethal 2, a dosage compensation gene of Drosophila, undergoes sex-specific regulation and encodes a protein with a RING finger and a metallothionein-like cysteine cluster. The EMBO journal, 14, 2884–2895.

27. Li, F., Schiemann, A.H. and Scott, M.J. (2008) Incorporation of the noncoding roX RNAs alters the chromatin-binding specificity of the Drosophila MSL1/MSL2 complex. Mol Cell Biol, 28, 1252–1264.

28. Morra, R., Yokoyama, R., Ling, H. and Lucchesi, J.C. (2011) Role of the ATPase/helicase maleless (MLE) in the assembly, targeting, spreading and function of the male-specific lethal (MSL) complex of Drosophila. Epigenetics Chromatin, 4, 6.

29. Kelley, R.L., Meller, V.H., Gordadze, P.R., Roman, G., Davis, R.L. and Kuroda, M.I. (1999) Epigenetic spreading of the Drosophila dosage compensation complex from roX RNA genes into flanking chromatin. Cell, 98, 513–522.

30. Villa, R., Jagtap, P.K.A., Thomae, A.W., Campos Sparr, A., Forne, I., Hennig, J., Straub, T. and Becker, P.B. (2021) Divergent evolution toward sex chromosome-specific gene regulation in Drosophila. Genes Dev.

31. Valsecchi, C.I.K., Basilicata, M.F., Georgiev, P., Gaub, A., Seyfferth, J., Kulkarni, T., Panhale, A., Semplicio, G., Manjunath, V., Holz, H. et al. (2021) RNA nucleation by MSL2 induces selective X chromosome compartmentalization. Nature, 589, 137–142.

32. Urban, J.A., Urban, J.M., Kuzu, G. and Larschan, E.N. (2017) The Drosophila CLAMP protein associates with diverse proteins on chromatin. PLoS One, 12, e0189772.

33. Rieder, L.E., Koreski, K.P., Boltz, K.A., Kuzu, G., Urban, J.A., Bowman, S.K., Zeidman, A., Jordan, W.T., 3rd, Tolstorukov, M.Y., Marzluff, W.F. et al. (2017) Histone locus regulation by the Drosophila dosage compensation adaptor protein CLAMP. Genes Dev, 31, 1494–1508.

34. Rieder, L.E., Jordan, W.T. and Larschan, E.N. (2019) Targeting of the Dosage-Compensated Male X-Chromosome during Early Drosophila Development. Cell Reports, 29, 4268-+.

35. Valsecchi, C.I.K., Basilicata, M.F., Semplicio, G., Georgiev, P., Gutierrez, N.M. and Akhtar, A. (2018) Facultative dosage compensation of developmental genes on autosomes in Drosophila and mouse embryonic stem cells. Nat Commun, 9, 3626.

36. Simpson, R.J., Yi Lee, S.H., Bartle, N., Sum, E.Y., Visvader, J.E., Matthews, J.M., Mackay, J.P. and Crossley, M. (2004) A classic zinc finger from friend of GATA mediates an interaction with the coiled-coil of transforming acidic coiled-coil 3. J Biol Chem, 279, 39789–39797.

37. Liew, C.K., Simpson, R.J.Y., Kwan, A.H.Y., Crofts, L.A., Loughlin, F.E., Matthews, J.M., Crossley, M. and Mackay, J.P. (2005) Zinc fingers as protein recognition motifs: Structural basis for the GATA-1/Friend of GATA interaction. P Natl Acad Sci USA, 102, 583–588.

38. Choi, S., Yamashita, E., Yasuhara, N., Song, J., Son, S.Y., Won, Y.H., Hong, H.R., Shin, Y.S., Sekimoto, T., Park, I.Y. et al. (2014) Structural basis for the selective nuclear import of the C2H2 zinc-finger protein Snail by importin beta. Acta Crystallogr D Biol Crystallogr, 70, 1050–1060.

39. Wang, B.S., Grant, R.A. and Pabo, C.O. (2001) Selected peptide extension contacts hydrophobic patch on neighboring zinc finger and mediates dimerization on DNA. Nat Struct Biol, 8, 589–593.

40. Dutnall, R.N., Neuhaus, D. and Rhodes, D. (1996) The solution structure of the first zinc finger domain of SWI5: a novel structural extension to a common fold. Structure, 4, 599–611.

41. Pavletich, N.P. and Pabo, C.O. (1993) Crystal structure of a five-finger GLI-DNA complex: new perspectives on zinc fingers. Science, 261, 1701–1707.

42. Buck-Koehntop, B.A., Stanfield, R.L., Ekiert, D.C., Martinez-Yamout, M.A., Dyson, H.J., Wilson, I.A. and Wright, P.E. (2012) Molecular basis for recognition of methylated and specific DNA sequences by the zinc finger protein Kaiso. Proc Natl Acad Sci U S A, 109, 15229–15234.

43. Bomar, M.G., Pai, M.T., Tzeng, S.R., Li, S.S. and Zhou, P. (2007) Structure of the ubiquitin-binding zinc finger domain of human DNA Y-polymerase eta. EMBO Rep, 8, 247–251.

44. Suzuki, N., Rohaim, A., Kato, R., Dikic, I., Wakatsuki, S. and Kawasaki, M. (2016) A novel mode of ubiquitin recognition by the ubiquitin-binding zinc finger domain of WRNIP1. FEBS J, 283, 2004–2017.

