## Supplementary Information for "Structural basis for interaction between CLAMP and MSL2 proteins involved in the specific recruitment of the dosage compensation complex in Drosophila"

***SUPPLEMENTARY METHODS***

### Protein expression and purification.

BL21 (DE3) cells were transformed with vectors encoding cDNAs of CLAMP derivatives and $MSL2^{618-655}$ fused with TEV-cleavable Thioredoxin and 6xHis-tag. Cells were grown in 700 ml of LB medium at 37°C to OD=0.6-1.0, then fresh 100 mM stock of Zinc Acetate was added to final concentration of 0.2 mM and protein expression was induced with 1 mM IPTG overnight at 18°C.

Cells were disrupted by sonication in 15 ml of buffer A (30 mM HEPES-KOH pH 7.5, 400 mM NaCl, 5 mM β-mercaptoethanol, 10 mM Imidazole, 0.1 % NP40, 5% (v/v) Glycerol, 0.1 mM $ZnCl_2$) containing 1 mM PMSF and Calbiochem Complete Protease Inhibitor Cocktail VII (1µL/1ml). After centrifugation lysate was applied to Ni-NTA column, and after washing with 20 ml of (30 mM HEPES-KOH pH 7.5, 400 mM NaCl, 5 mM β-mercaptoethanol, 30 mM Imidazole) protein was eluted with 15 ml of (30 mM HEPES-KOH pH 7.5, 400 mM NaCl, 5 mM β-mercaptoethanol, 300 mM Imidazole).

For cleavage of Thioredoxin-6xHis-tag sodium citrate was added to final concentration of 5mM, $ZnCl_2$ to 0.01 mM and 6x-His-tagged TEV protease was added at molar ratio approximately 1:50 directly to the eluted protein, mixture was incubated for 2 hours at room temperature and dialyzed overnight at 4°C against degassed 30 mM HEPES-KOH pH 7.5, 400 mM NaCl, 1 mM β-mercaptoethanol, 10 mM Imidazole, 0.1 mM $ZnCl_2$, then filtered and applied to Ni-NTA column, flowthrough was collected, dialyzed against degassed 20 mM Tris-HCl, pH 7.4, 0.1 mM $ZnCl_2$, 1 mM DTT and further purified using SOURCE15Q 4.6/100 column (GE Healthcare). Proteins were either eluted with 0-500 mM NaCl gradient (CLAMP deletion derivatives), or collected as flowthrough ($MSL2^{618-655}$). Sample homogeneity was confirmed with size-exclusion chromatography which was performed using Superdex 200 10/300GL column (GE Healthcare) in 20 mM Tris-HCl, pH 7.4, 150 mM NaCl, 0.1 mM $ZnCl_2$, 1 mM DTT.

### Fly crosses and transgenic lines

*Drosophila* strains were grown at 25°C under standard culture conditions. The transgenic constructs (Ubi-clamp or Ubi-MSL2) were injected into preblastoderm embryos using the φC31-mediated site-specific integration system at locus 86Fb (1). The emerging adults were crossed with the *y ac* $w^{1118}$ flies, and the progeny carrying the transgene in the 86Fb region were identified by $y^+$ pigmented cuticle. To assess the viability of transgenic lines expressing different CLAMP variants (CLAMP*), virgin $clamp^2$/CyO, GFP; Ubi:clamp*-HA/Ubi:clamp*-HA females were crossed with $clamp^2$/CyO, GFP; Ubi:clamp*-HA/Ubi:clamp*-HA males. Viability of transgenic flies expressing CLAMP* was calculated as the ratio of the homozygous males or females ($clamp^2/clamp^2$ ; Ubi:clamp*-HA/Ubi:clamp*-HA) relative to heterozygous male or females ($clamp^2$/CyO; Ubi:clamp*-HA/Ubi:clamp*-HA) divided by two. To assess the *in vivo* role of mutations in MSL2, the viability of females homozygous for the Ubi:msl2*-FLAG transgene was assessed in homozygous transgenic lines (Ubi:msl2*-FLAG/Ubi:msl2*-FLAG), where msl2* expresses WT or one of the mutant variants of MSL2. Fly protein extracts were performed as described (2).

### Polytene chromosome staining

*Drosophila* 3[rd] instar larvae were cultured at 18°C under standard conditions. Immunostaining of polytene chromosomes was performed as described (3). The following primary antibodies were used: rabbit anti-MSl1 at 1:500 dilution, rabbit anti-Msl2 at 1:500 dilution, and monoclonal mouse anti-FLAG at 1:50 dilution. The secondary antibodies were Alexa Fluor 488 goat anti-mouse 1:2000 and Alexa Fluor 555 goat anti-rabbit 1:2000 (Invitrogen). The polytene chromosomes were co-stained with DAPI. Images were acquired on the Nikon Eclipse T*i* fluorescent microscope using Nikon DS-Qi2 digital camera, processed with ImageJ 1.50c4 and Fiji bundle 2.0.0-rc-46. 3-4 independent stainings and 4-5 samples of polytene chromosomes were performed with each transgenic line.

### Analysis of C2H2 zinc-fingers amino-acid composition

We compared amino acid residues of the CLAMP N-terminal zinc finger at DNA-binding positions with their average abundance in C2H2 zinc-fingers. We developed a hidden Markov model of the C2H2 domain sequence based on 109192 representative domains from the Pfam database (4). We used it to calculate each residue's probabilities at given positions (see Supplementary Figure S7). The probability of each residue in a completely random sequence would be 5%. The CLAMP N-terminal zinc finger has histidine at -1 position, leucines at +1 and +3, alanines at +2 and +6 (+2 and +6 positions are not conserved in some species, substituted for threonine or histidine, respectively). Histidine at -1 position is found here in 4.8% of zinc fingers according to our model; leucine at +1 is rarely used for DNA recognition (5) but is present at this position in 7.2% of zinc fingers. Alanine or threonine at +2 is found in 11.2% zinc-fingers. Leucine is rarely found at position +3 (3.3% of all zinc fingers), a polar residue in most (93.3%) zinc fingers. Residue at +6 position also most often is polar (78.7%), but alanines (and histidines, which are found here in some species) sometimes (6.1% altogether) are present at this position (6,7). The first residue of the second beta-sheet often makes non-specific contacts with backbone phosphate, and arginine or lysine is usually (75.6%) found at this position (8), the CLAMP N-terminal zinc-finger has nontypical hydrophobic leucine at this position, which is found here only in 0.9% of zinc fingers. De novo prediction of the target site (9) of the CLAMP N-terminal zinc finger yields low-score, ambiguous results (the position-weight matrix is shown in Supplementary Figure S8). Altogether, these analyses suggest that the CLAMP N-terminal zinc-finger is very uncommon as a DNA-binding zinc finger and is most likely not involved in DNA-binding.

**NMR Spectroscopy**

The standard triple resonance experiments were performed for the sequential assignments of the backbone $^{1}$H, $^{13}$C and $^{15}$N nuclei of both, CLAMP$^{87\text{-}153}$ and MSL2$^{618\text{-}655}$ protein fragments: HNCO, HN(CA)CO, CBCA(CO)NH, and HNCACB. Additionally, for the side-chain assignment and structure calculation, the $^{13}$C-HCCH-TOCSY, DQF-COSY, $^{13}$C-$^{1}$H HSQC-NOESY, and $^{15}$N-$^{1}$H HSQC-NOESY spectra were collected for CLAMP$^{87\text{-}153}$.

The acquired data were processed using NMRPipe (10), and analyzed using NMRFAM-Sparky software (11).

**NMR chemical shifts assignments and data deposition**

Backbone $^{1}$H, $^{13}$C and $^{15}$N resonance assignment was performed manually using NMRFAM-Sparky (11). Backbone amide $^{1}$H and $^{15}$N resonance assignments of CLAMP$^{87\text{-}153}$ and MSL2$^{618\text{-}655}$ were achieved for all non-proline residues. Side-chain signals of CLAMP$^{87\text{-}153}$ were assigned manually using the information on the backbone assignments. The $^{1}$H, $^{15}$N and $^{13}$C chemical shifts of CLAMP$^{87\text{-}153}$ have been deposited into the BioMagResBank (www.bmrb.wisc.edu) under the accession number BMRB-34600.

**Protein chain mobility, secondary structure elements and relaxation**

The protein chain mobility and secondary structure elements of CLAMP and MSL2 were identified with TALOS+ program using $^{13}$Cα, $^{13}$Cβ, $^{13}$C′ and $^{15}$N chemical shifts (12). These data were also used to get backbone torsion angle restraints for structure calculations.

Correlation time of protein tumbling was calculated from the NMR relaxation data. The obtained values were used to estimate protein radius using the Stokes' law:

$$\tau_c = \frac{4\pi\eta r^3}{3kT}$$

Estimation was done with the value 8.90×10$^{-4}$ Pa for the water viscosity at 25 °C (13)

**NMR Restraints generation and Structure calculation**

The solution structure of CLAMP$^{87\text{-}153}$ was calculated using following set of restraints: (1) distance restraints obtained from $^{13}$C-$^{1}$H HSQC-NOESY and $^{15}$N-$^{1}$H HSQC-NOESY spectra; (2) torsion angle restraints obtained from chemical shifts using TALOS+ program; (3) H/D exchange rates (Supplementary Table S1A).

The structure calculation was performed using ARIA (14) and CNS (15) programs. Structural statistics is shown at Supplementary Table S1A and B. Ramachandran plot (Supplementary Figure S16) shows that all residues conformations are correct.

CLAMP N-terminal C2H2 domain is a classic example of zinc-finger in terms of both fold and metal-binding environment (Figure 1A). Its 26-residue sequence adopts a ββα tertiary fold consisting of short β-hairpin ($\beta_1$:F127-C129/$\beta_2$:S133-F136) followed by long C-terminal α-helix ($\alpha_3$:L139-T150) which is in agreement with the secondary structure calculation based on chemical shifts. The fold is further stabilized by a hydrophobic core that is formed around the side-chain of L142 residue. The metal coordination site consists of two cysteine and two histidine residues ($C_2H_2$).

The β-hairpin is supported by strong H-bond between carbonyl group of F127 and $H^N$ atom of F136, as was shown by H/D exchange experiments (Supplementary Figure S2). Apart from the F136 $H^N$, there are also two $H^N$ atoms possessing much longer exchange rate than the others, namely V131 and C132. The last two atoms are not participating in H-bonds, but are placed near the $Zn^{2+}$ ion. Upstream of Zinc-finger, CLAMP is unstructured, and the presence of upstream fragment has no impact on Zinc-finger structure. It was demonstrated by expression of $^{15}$N-labelled CLAMP fragments: CLAMP$^{1\text{-}153}$, CLAMP$^{40\text{-}153}$, as well as CLAMP$^{1\text{-}117}$, with the subsequent HSQC spectra measurement (Supplementary Figure S3).

Molecular modeling of MSL2$^{618\text{-}655}$ structure was performed based on chemical shifts using CS-Rossetta (16). The modelling suggests a possibility of β-hairpin formation at V634−N638 and G641−N647. At the same time, the order parameter S2 for these residues does not exceed 0.7 (Supplementary Figure S10), which corresponds to unstructured protein chain. To validate the formation of β-hairpins in MSL2$^{618\text{-}655}$ we assigned Hα-atoms using HNHA and HBHA(CO)NH spectra and measured 3D 15N-1H HSQC-NOESY spectra, but no NOE between V634−N638 and G641−N647 were found. Thus, we approved the absence of β-hairpins in the corresponding region.

**Chemical Shifts perturbation**

Interaction of CLAMP with MSL2 was studied using NMR titration experiments. $^{15}$N-labelled MSL2$^{618\text{-}655}$ at a concentration of 150 μM was used. The unlabelled CLAMP$^{87\text{-}153}$ concentration increased from 1:1 to 1:13 protein–protein ratio. For each titration point, a $^{15}$N–$^{1}$H SOFAST HMQC spectrum (17) was recorded.

The titration of $^{15}$N-labeled CLAMP by MSL2 were performed using CLAMP$^{87\text{-}153}$, CLAMP$^{40\text{-}153}$, and CLAMP$^{1\text{-}153}$ constructs and unlabelled MSL2$^{618\text{-}655}$.

The chemical shift perturbation data were calculated using the formula: $((\Delta\delta(^1H^i))^2 + (\Delta\delta(^{15}N^i)/25)^2)^{½}$.

The value of $K_d$ were estimated from NMR titration experiments carried out at 25 °C. 26 $^1$H amide resonances were used in non-linear fitting of $K_d$ values by the following equation (18):

$$\Delta\delta_{obs} = \frac{\Delta\delta_{max}}{2[P]_0}\left[(K_d + [P]_0 + [L]_0) - \sqrt{(K_d + [P]_0 + [L]_0)^2 - 4[P]_0[L]_0}\right]$$

where $P_0$ and $L_0$ are the total concentrations of CLAMP and the MSL2 in each titration step, $\Delta\delta_{obs}$ is the change of chemical shift value, and $\Delta\delta_{max}$ is the maximum change of the chemical shift accepted by the difference between the signal in free protein and protein in the presence its partner in the maximum concentration.

*SUPPLEMENTARY TABLES*

**Supplementary Table S1.** Statistics for the ensemble of the calculated 20 structures of the CLAMP$^{87\text{-}153}$. No NOE or dihedral angle violations are above 0.5 Å and 5° respectively.

A. Restraints used in the structure calculation

| | | | |
|---|---|---|---|
| Total NOEs | 355 | Total dihedral angles | 40 |
| Long range ($\|i - j\| > 4$) | 30 | Phi (ϕ) | 20 |
| Medium ($1 < \|i - j\| \leq 4$) | 43 | Psi (ψ) | 20 |
| Sequential ($\|i - j\| = 1$) | 91 | H-bonds | 1 |
| Intraresidue | 191 | | |

B. Restraint violations and structural statistics (for 20 structures)

| Average RMSD | ensemble of 20 final structures | representative structure |
|---|---|---|
| From experimental restraints | | |
| Distance (Å) | 0,023±0.002 | 0,026 |
| Dihedral (°) | 0,57±0,12 | 0,49 |
| From idealized covalent geometry | | |
| Bonds (Å) | 0,0018±0,0001 | 0,002 |
| Angles (°) | 0,397±0,014 | 0,414 |
| Impropers (°) | 0,295±0,014 | 0,272 |
| Ramachandran plot statistics | | |
| % of residues in most favorable region of Ramachandran plot | 100 | 100 |
| % of residues in disallowed region of Ramachandran plot | 0 | 0 |

C. Superimposition on the representative structure (Å)

| | |
|---|---|
| Backbone (C, Cα, N) RMSD over the structured protein core (residues 126-150) | 0.46±0,10 |

**Supplementary Table S2. (A)** Comparison of the viability of males and females upon rescue of the clamp$^2$/clamp$^2$ and clamp$^2$/+ mutant background with CLAMP proteins expressed in transgenic constructs. **(B)** Comparison of the viability of males and females upon ectopic expression of MSL2 proteins shown as triplicate experiments.

**(A)**

| | clamp$^2$/+ | | clamp$^2$/clamp$^2$ | |
|---|---|---|---|---|
| | ♂ | ♀ | ♂ | ♀ |
| clamp $^{WT}$ | 389 | 363 | 104 | 102 |
| clamp $^{K146E}$ | 199 | 192 | 5 | 6 |
| clamp $^{L139A\ K146E}$ | 203 | 215 | 6 | 6 |
| clamp $^{K146E\ R147E}$ | 105 | 100 | 7 | 7 |

**(B)**

| | ♀ | ♂ |
|---|---|---|
| msl2 Y643A | 56 | 64 |
| | 49 | 56 |
| | 84 | 93 |
| msl2 E639A Y643A | 80 | 78 |
| | 91 | 88 |
| | 82 | 91 |
| msl2 L633A V634AvV E639A Y643A | 107 | 106 |
| | 87 | 87 |
| | 82 | 77 |

**Supplementary Table S3.** Oligonucleotides used for cloning. Restriction enzyme sites are shown in small letters, the corresponding enzymes are noted. Nucleotide substitutions in mutagenic primers are also shown in small letters.

| | Direct | Reverse | REs |
|---|---|---|---|
| CLAMP 87-153 | CTGgaattcATGGAAGACCTTACCAA | AACgtcgacTTCCCCGTCTGTATGCAT | *Hinc*II,*Sal*I |
| CLAMP 40-153 | CTGgaattcATGAAAACGGAGCAGCAGC | AACgtcgacTTCCCCGTCTGTATGCAT | *Eco*RI,*Sal*I |
| MSL2 618-655 | TCTggatccATAAGCCTAGTGCCGC | GTGgaattcCTAATCAAGGGGCT | *Bam*HI, *Eco*RI |
| MSL2 | CGAGTACTGGTGGCGTGGGTCGGACCGAC | TCTgtcgacCAAGTCATCCGAGCCCGACA | *Sca*I, *Sal*I |
| CLAMP 1-153 | CTGgaattcATGGAAGACCTTACCAA | AACgtcgacTTCCCCGTCTGTATGCAT | *Eco*RI,*Sal*I |
| CLAMP 1-153 $^{I122A}$ | CTAACACCgcATCCAACATAAGC | CTTATGTTGGATgcGGTGTTAGC | |
| CLAMP 1-153 $^{H138A}$ | GTTCCCTgcTTTGGCACTTC | GAAGTGCCAAAgcAGGGAACAT | |
| CLAMP 1-153 $^{L139A}$ | CCCTCATgcGGCACTTCTTAATG | CATTAAGAAGTGCCgcATGAGGGAAC | |
| CLAMP 1-153 $^{L141A}$ | CATTTGGCAgcTCTTAATGCTC | GAGCATTAAGAgcTGCCAAATG | |
| CLAMP 1-153 $^{N143A}$ | CACTTCTTgcTGCTCATGAGGAG | CTCATGAGCAgcAAGAAGTGCC | |
| CLAMP 1-153 $^{K146E}$ | CTCATgAGCGGATGCATACAGAC | GTATGCATCCGCTcATGAGCATTAAG | |
| CLAMP 1-153 $^{R147E}$ | GCTCATAAGgaGATGCATACAGACG | GTCTGTATGCATCtcCTTATGAGC | |
| CLAMP 1-153 $^{K146ER147E}$ | GTCCATgAGgaGATGCATACAGACG | GTCTGTATGCATCtcCTCATGAGC | |
| CLAMP 1-153 $^{N143AK146ER147E}$ | GCTCATgAGgaGATGCATACAGACG | CTCATGAGCAgcAAGAAGTGCC | |
| HA-CLAMP | CTGgaattcATGGAAGACCTTACCAA | TAGgtcgacCTATAACCCACCGATAATC | *Eco*RI,*Sal*I |
| MSL2-FLAG | TTCgatatcATGGCCCAGACGGCATACTT | TCTcccgggCAAGTCATCCGAGCCCGA | *Eco*RV,*Sma*I |
| MSL2-FLAG $^{Y643A}$ | AAGGCGAGgcCCAGGGCTTCAATATCTT | AGCCCTGGgcCTCGCCTTTCTCATTCTG | |
| MSL2-FLAG $^{E639AY643A}$ | TCAGAATGcGAAAGGCGAGgcCCAGG | CGCCTTTCgCATTCTGAACAAGCACC | |
| MSL2-FLAG $^{L633AV634A}$ $^{E639AY643A}$ | CATCCTgcGGcGCTTGTTCAGAAT | AACAAGCgCCgcAGGATGCTGGGA | |
| amMSL2 | CAAggatccATGAATGCCACAAGTCTTTACG | CTTgtcgacTCACACATCAATTTGTATATCACTGTC | *Bam*HI, *Sal*I |
| amCLAMP 1-204 | GACggatccATGGTCAAAGGCAACACATC | TTGgtcgacTTATTGAAGATTACTAGGTGTTGTATTATG | *Bam*HI, *Sal*I |
| amMSL2 338-432 | GGTggatccATTCGACCACATATACCTGAATTAC | CTTgtcgacTCACACATCAATTTGTATATCACTGTC | *Bam*HI, *Sal*I |

***SUPPLEMENTARY FIGURES***

**Supplementary Figure S1.** Protein chain mobility and secondary structure of CLAMP$^{87-153}$. RCI is Random Coil Index, values of model-free order parameters ($S^2$) are shown.

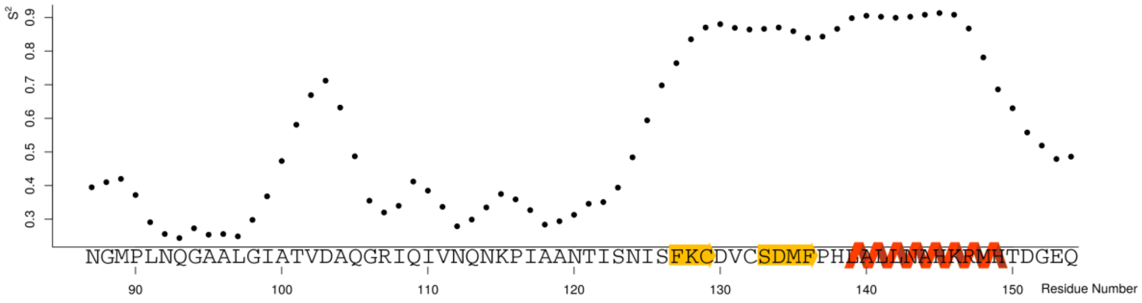

**Supplementary Figure S2.** H/D exchange experiments show much slower exchange rate for V131, C132, and F136 compared to the other residues.

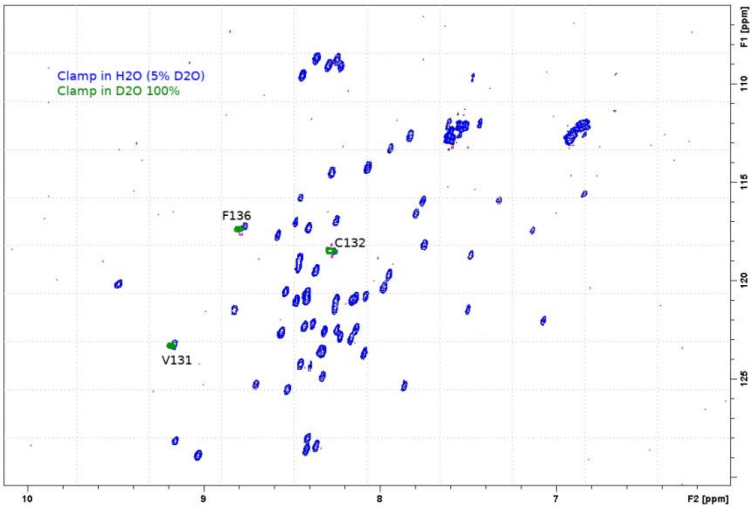

**Supplementary Figure S3. (A)** CLAMP$^{40\text{-}153}$ vs CLAMP$^{87\text{-}153}$ chemical shifts. Partial amino acid assignment (Performed for residues 87-153) is shown. **(B)** Overlay of CLAMP$^{1\text{-}153}$ over CLAMP$^{40\text{-}153}$ chemical shifts. **(C)** CLAMP$^{1\text{-}117}$ vs CLAMP$^{1\text{-}153}$ chemical shifts.

(A)

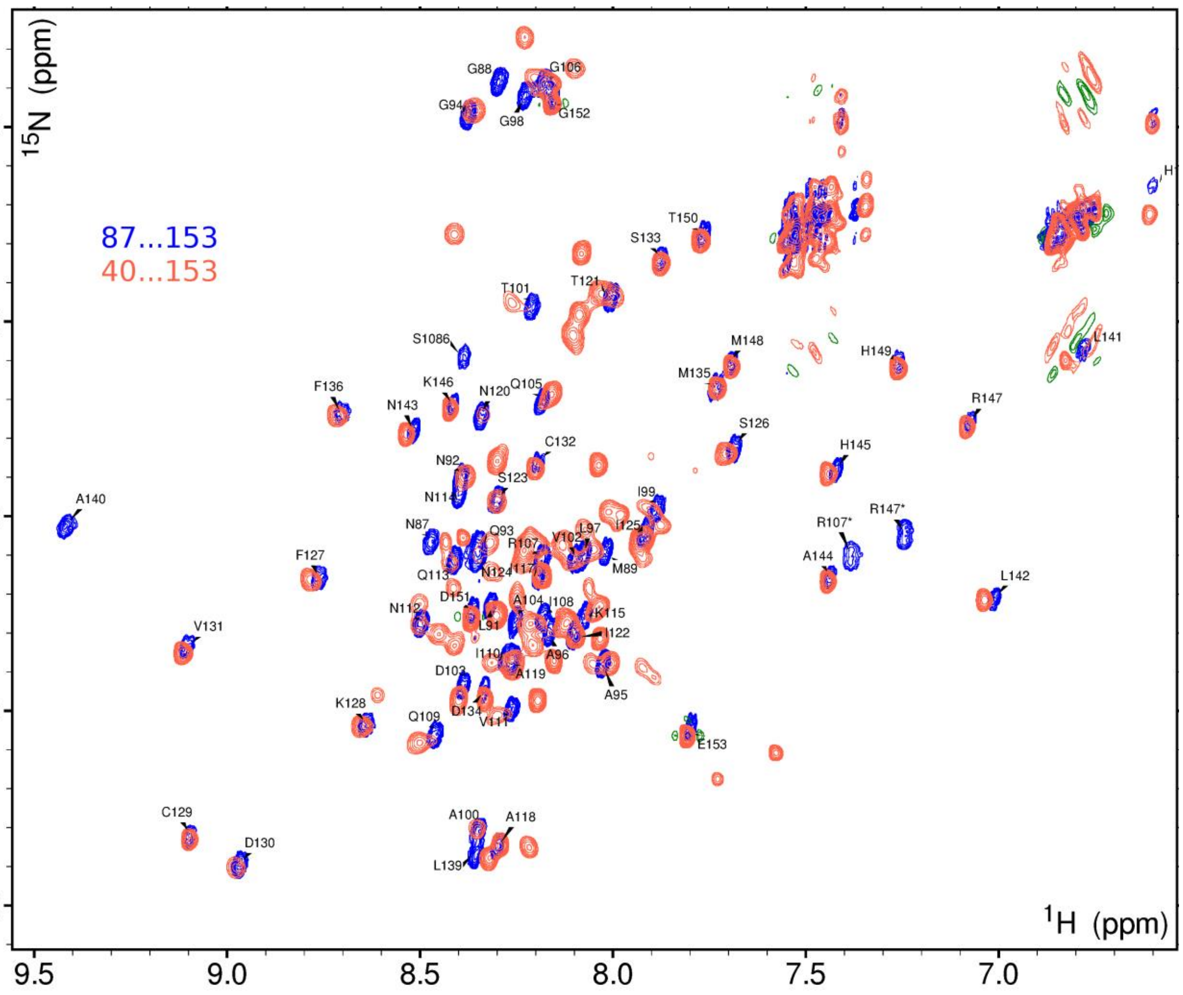

(B)

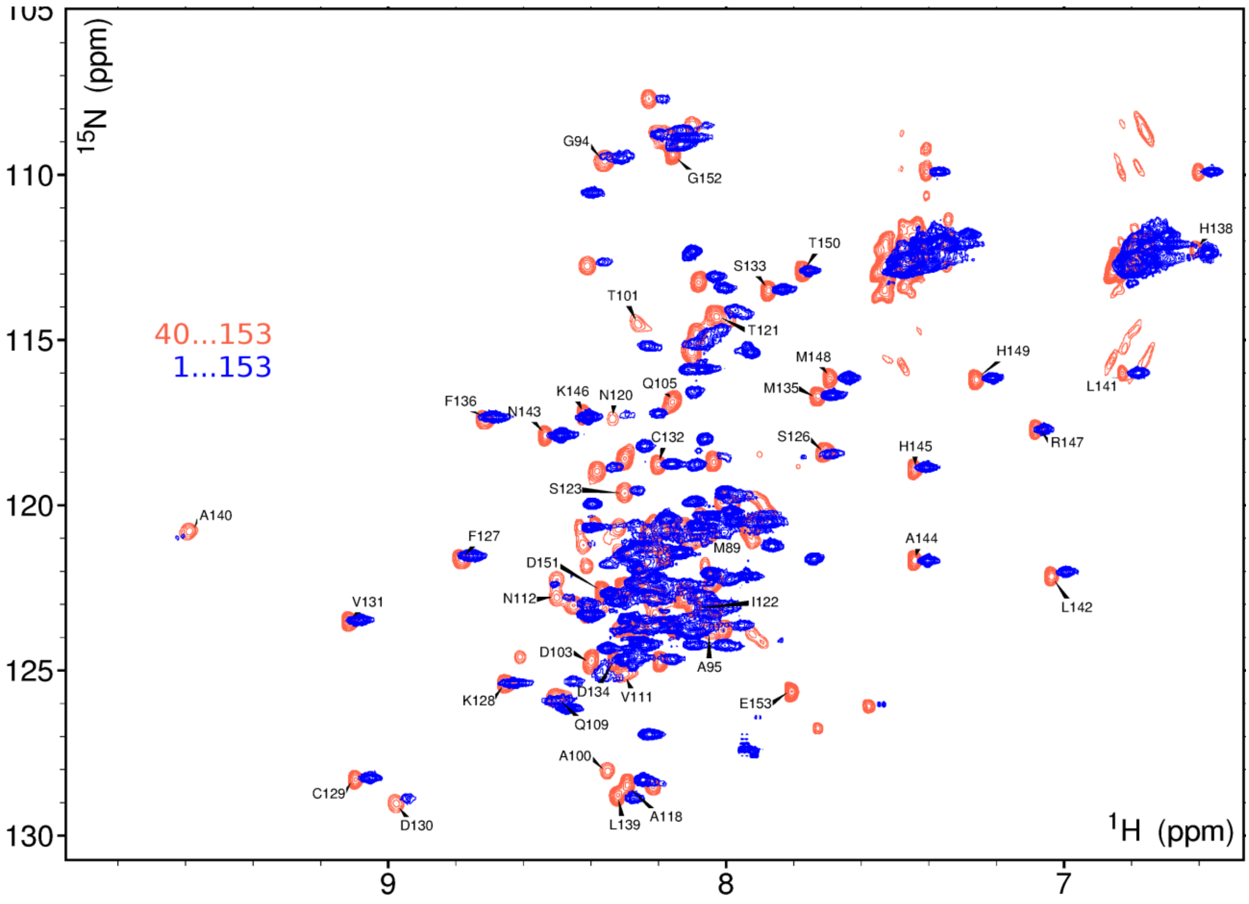

40...153
1...153
$^{15}$N (ppm)
$^{1}$H (ppm)
105
110
115
120
125
130
9
8
7
G94
G152
T150
S133
H138
T101
T121
M148
H149
L141
M135
F136
N143
K146
N120
Q105
C132
S126
H145
R147
S123
A140
F127
M89
A144
D151
N112
I122
L142
V131
D103
A95
D134
V111
E153
K128
Q109
A100
C129
L139
A118
D130

(C)

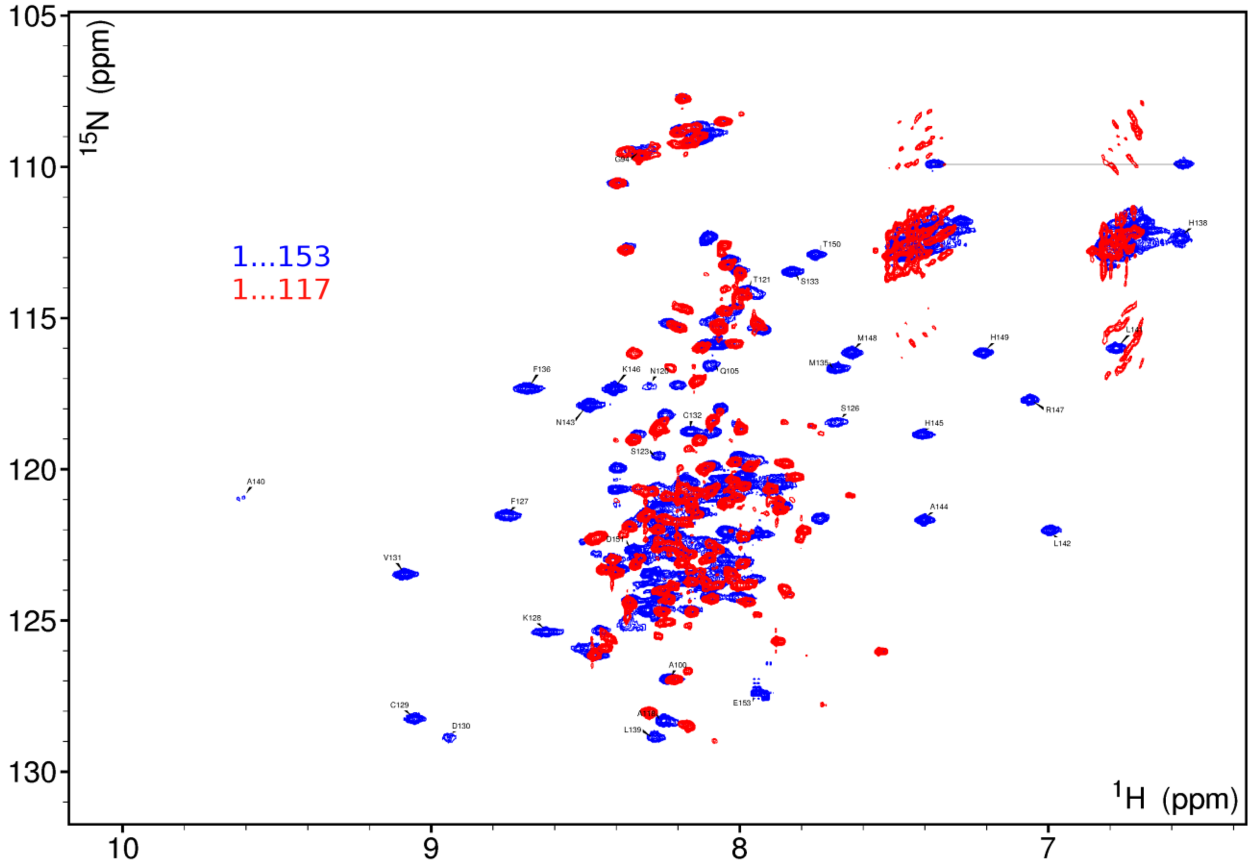

**Supplementary Figure S4.** Multiple sequence alignment of MSL2-interacting CLAMP N-terminal region from various insects. Position of zinc-finger domain is shown. Phylogenetic positions of taxa are shown according to (19).

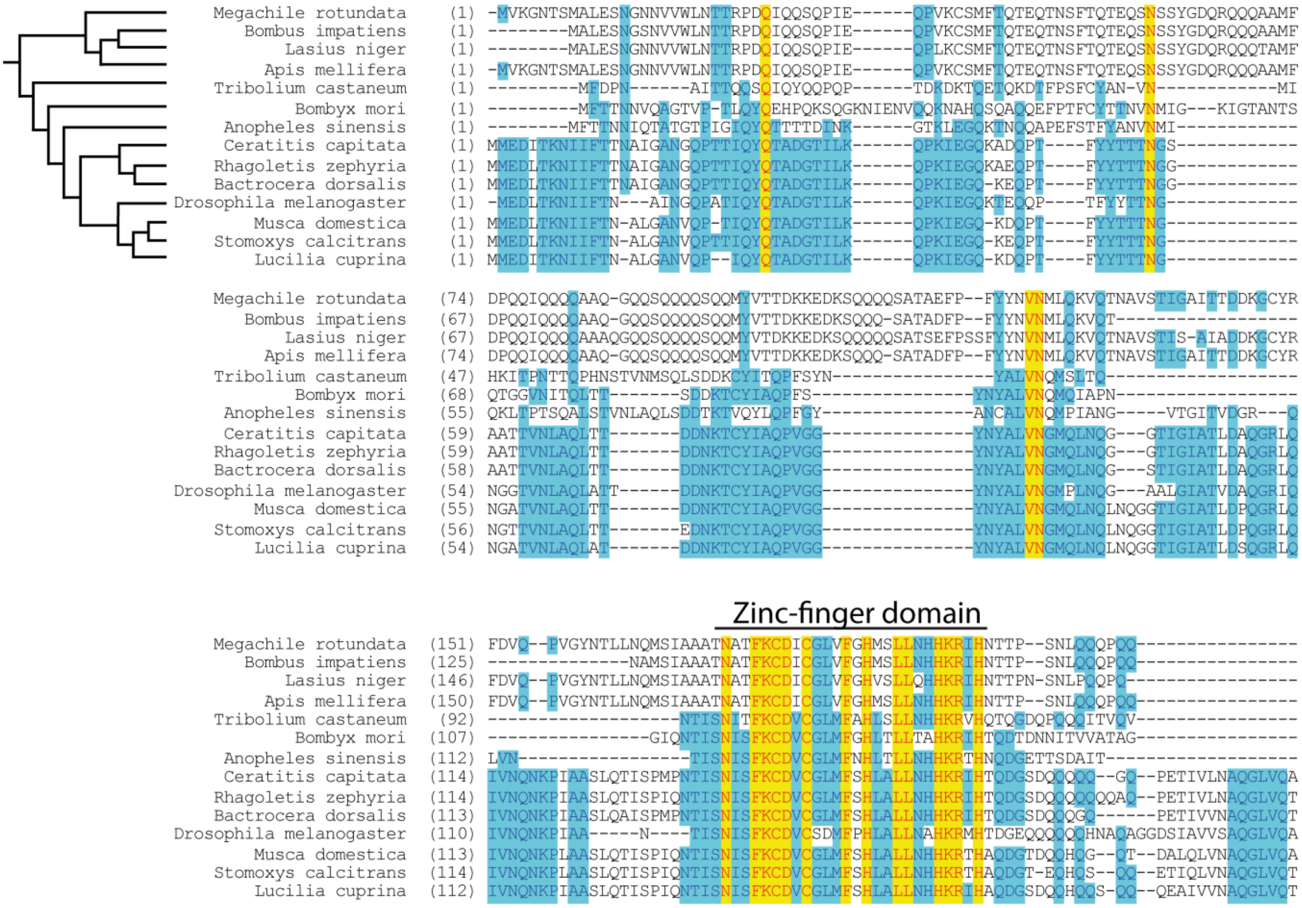

**Supplementary Figure S5.** Plots of the relaxation time for the $^{15}N$ nuclei of the backbone amide groups of $CLAMP^{40\text{-}153}$ and $CLAMP^{87\text{-}153}$ as a function of residue number. Shown are longitudinal relaxation time $T_1$, s (upper plot) and transverse relaxation time $T_2$, s (lower plot) measured at 700 MHz and 25 ºC. Correlation time of protein tumbling ($\tau_c$) was calculated from the NMR relaxation data for $CLAMP^{87\text{-}153}$ (2.7 ns) and $CLAMP^{40\text{-}153}$ (2.8 ns). Thus, the presence of residues 40-86 does not affect significantly $\tau_c$. The obtained values were used to estimate protein radius using the Stoke's law and gave ~1.4 nm. It is almost the same value as calculated for Zinc-finger alone (~1.1 nm for the residues 126-150).

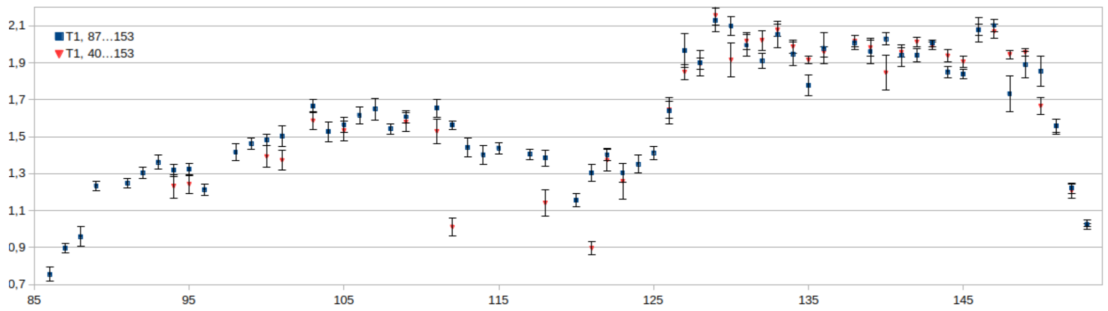


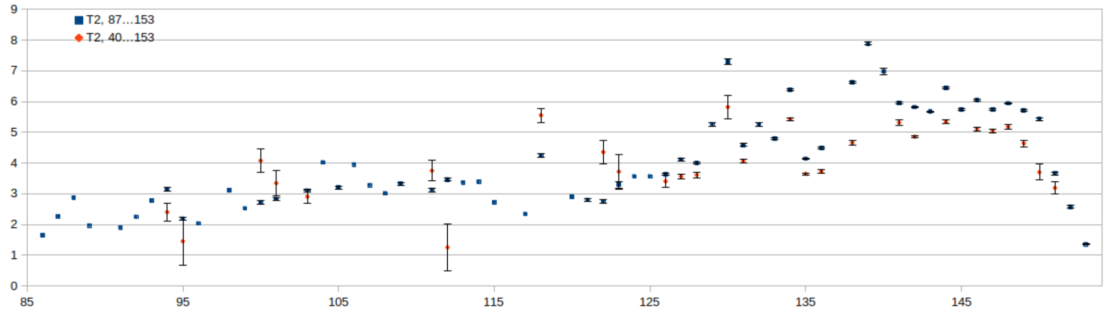

**Supplementary Figure S6.** $^{15}$N-HSQC Spectra of $^{15}$N-labelled CLAMP$^{40\text{-}153}$ titrated with increasing concentrations of unlabeled MSL2$^{618\text{-}655}$ peptide.

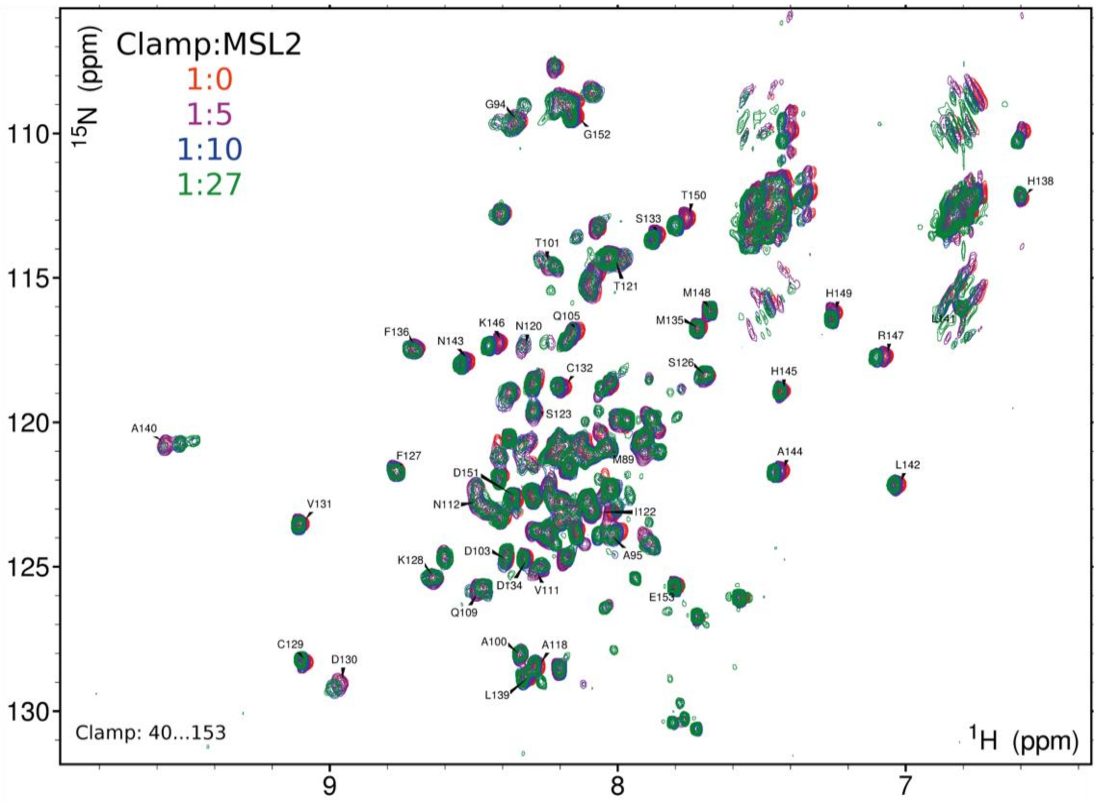

**Supplementary Figure S7.** Analysis of global C2H2 fingers amino-acid composition. **(A)** A logo representing sequence alignment and profile hidden Markov model for 109192 C2H2 zinc-fingers calculated with Skylign (20). **(B)** Probabilities of residues at known DNA-interacting positions are shown below.

A

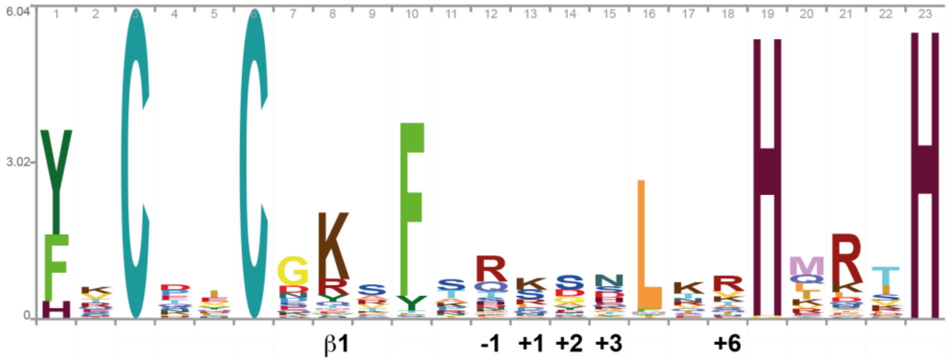


B

**-1**

| Residue | Probability | Residue | Probability | Residue | Probability | Residue | Probability |
|---|---|---|---|---|---|---|---|
| R | 0.393 | D | 0.043 | E | 0.020 | I | 0.006 |
| Q | 0.156 | N | 0.040 | V | 0.018 | W | 0.005 |
| T | 0.091 | K | 0.033 | L | 0.017 | M | 0.005 |
| S | 0.057 | A | 0.026 | F | 0.009 | C | 0.004 |
| H | 0.048 | Y | 0.021 | G | 0.007 | P | 0.001 |

**+1**

| Residue | Probability | Residue | Probability | Residue | Probability | Residue | Probability |
|---|---|---|---|---|---|---|---|
| K | 0.259 | Q | 0.062 | H | 0.011 | Y | 0.006 |
| S | 0.224 | N | 0.050 | V | 0.008 | F | 0.006 |
| R | 0.110 | A | 0.039 | M | 0.008 | D | 0.004 |
| L | 0.072 | T | 0.034 | G | 0.007 | W | 0.004 |
| P | 0.072 | E | 0.012 | I | 0.007 | C | 0.004 |

**+2**

| Residue | Probability | Residue | Probability | Residue | Probability | Residue | Probability |
|---|---|---|---|---|---|---|---|
| S | 0.312 | E | 0.043 | Q | 0.016 | P | 0.008 |
| D | 0.153 | H | 0.036 | F | 0.015 | I | 0.007 |
| G | 0.138 | T | 0.034 | K | 0.013 | L | 0.006 |
| A | 0.075 | N | 0.020 | C | 0.013 | W | 0.006 |
| Y | 0.073 | R | 0.020 | V | 0.010 | M | 0.003 |

**+3**

| Residue | Probability | Residue | Probability | Residue | Probability | Residue | Probability |
|---|---|---|---|---|---|---|---|
| N | 0.248 | Q | 0.054 | Y | 0.028 | I | 0.005 |
| S | 0.137 | K | 0.054 | G | 0.023 | F | 0.005 |
| D | 0.109 | A | 0.051 | M | 0.014 | C | 0.004 |
| H | 0.103 | T | 0.049 | R | 0.014 | W | 0.002 |
| E | 0.055 | L | 0.033 | V | 0.008 | P | 0.002 |

**+6**

| Residue | Probability | Residue | Probability | Residue | Probability | Residue | Probability |
|---|---|---|---|---|---|---|---|
| R | 0.332 | S | 0.051 | L | 0.023 | H | 0.007 |
| V | 0.132 | Q | 0.051 | D | 0.013 | C | 0.002 |
| K | 0.109 | I | 0.045 | Y | 0.010 | W | 0.002 |
| T | 0.074 | N | 0.037 | M | 0.009 | F | 0.002 |
| A | 0.061 | E | 0.029 | G | 0.009 | P | 0.001 |

**β1**

| Residue | Probability | Residue | Probability | Residue | Probability | Residue | Probability |
|---|---|---|---|---|---|---|---|
| K | 0.613 | M | 0.017 | F | 0.008 | N | 0.005 |
| R | 0.143 | A | 0.015 | S | 0.007 | P | 0.004 |
| Y | 0.069 | E | 0.015 | H | 0.007 | C | 0.003 |
| Q | 0.036 | W | 0.012 | V | 0.006 | G | 0.002 |
| T | 0.022 | L | 0.009 | I | 0.006 | D | 0.002 |

**Supplementary Figure S8.** De novo prediction of possible DNA-binding by CLAMP N-terminal zinc-finger. **(A)** Position-weight matrix. **(B)** Schematic representation of DNA-binding site.

**A**

| base | 1 | 2 | 3 |
|---|---|---|---|
| a | 0.251 | 0.214 | 0.116 |
| c | 0.155 | 0.121 | 0.376 |
| g | 0.255 | 0.236 | 0.242 |
| t | 0.340 | 0.428 | 0.266 |

**B**

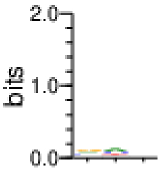

**Supplementary Figure S9.** Testing of the impact of point mutations on CLAMP-MSL2 interaction using yeast two-hybrid assay. Growth assay plates without histidine are shown (yeasts are unable to grow on this medium in the absence of interaction). AD stands for Activation Domain, BD – for DNA-Binding Domain of GAL4 protein.

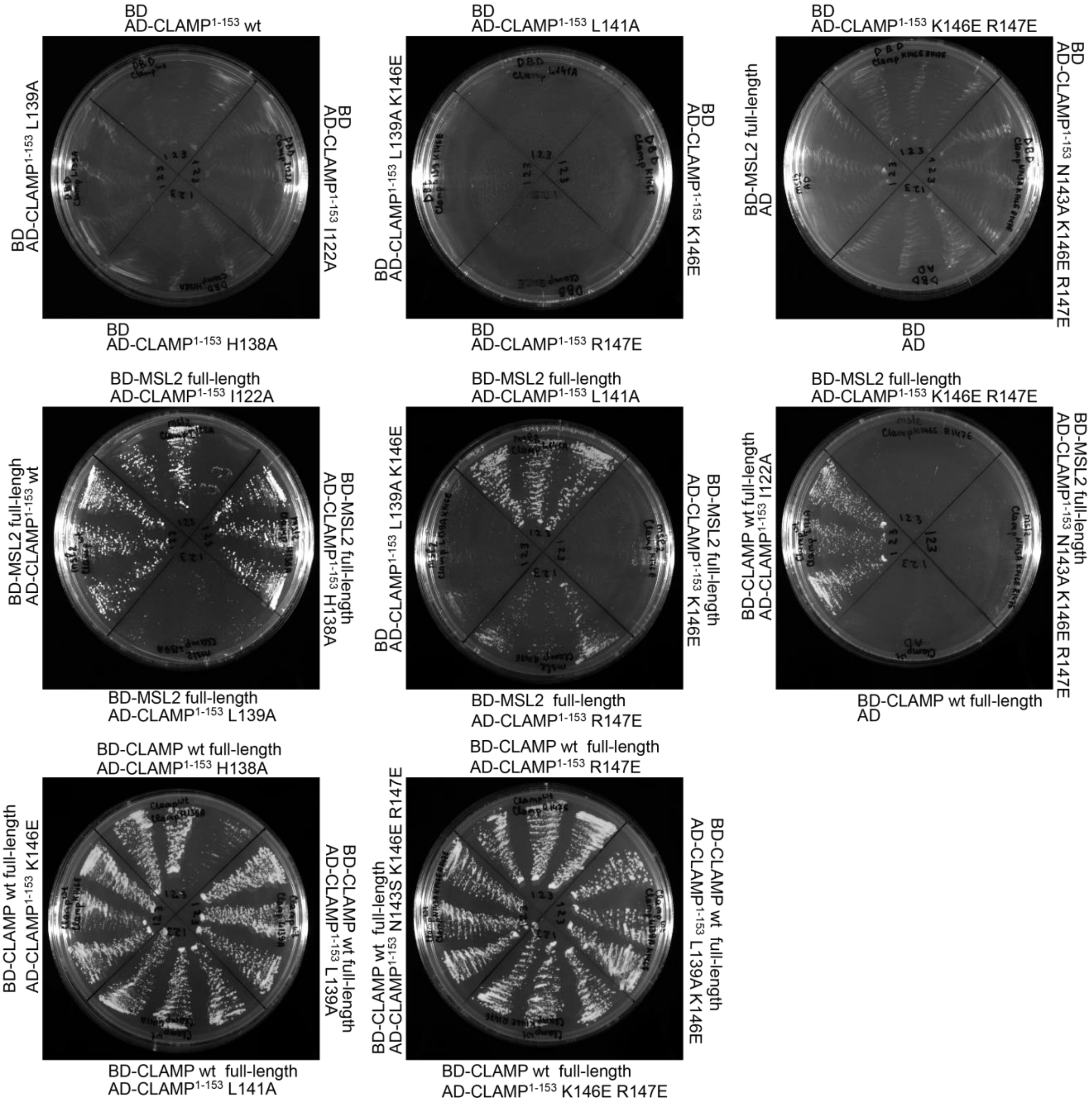

**Supplementary Figure S10.** Protein chain mobility of MSL2[618-655]. RCI is Random Coil Index, values of model-free order parameters ($S^2$) are shown.

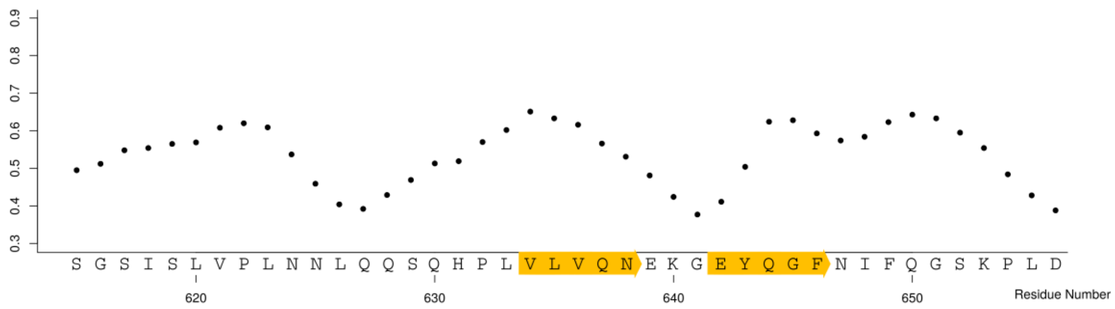

**Supplementary Figure S11.** $^{15}$N-HSQC Spectra of $^{15}$N-labelled MSL2$^{618-655}$ titrated with increasing concentrations of unlabeled CLAMP$^{40-153}$ peptide.

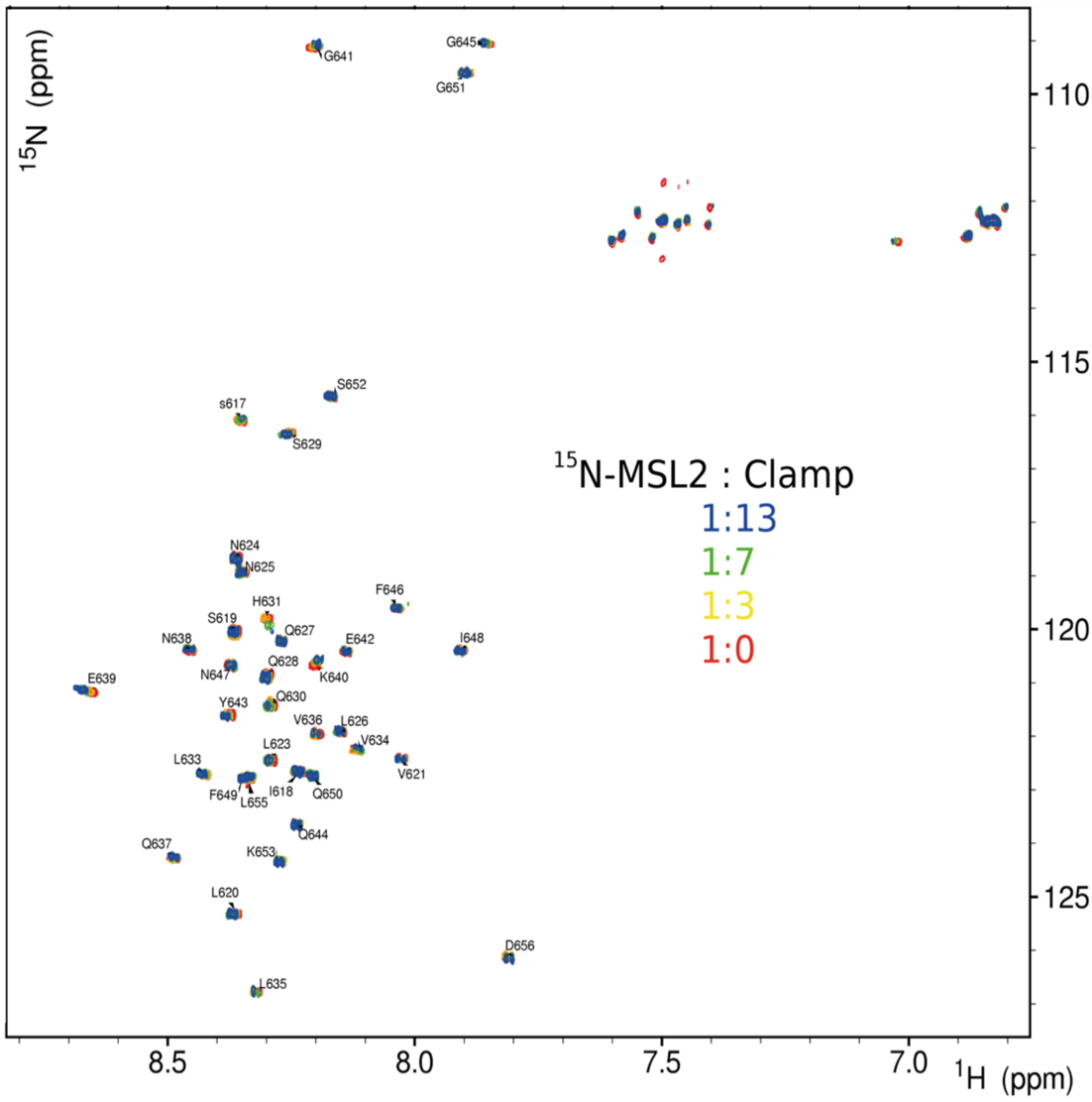

**Supplementary Figure S12.** Multiple sequence alignment of MSL2 CXC and C-terminal domains from various insects. CLAMP-interacting residues are shown in bold at the *D. melanogaster* MSL2 sequence.

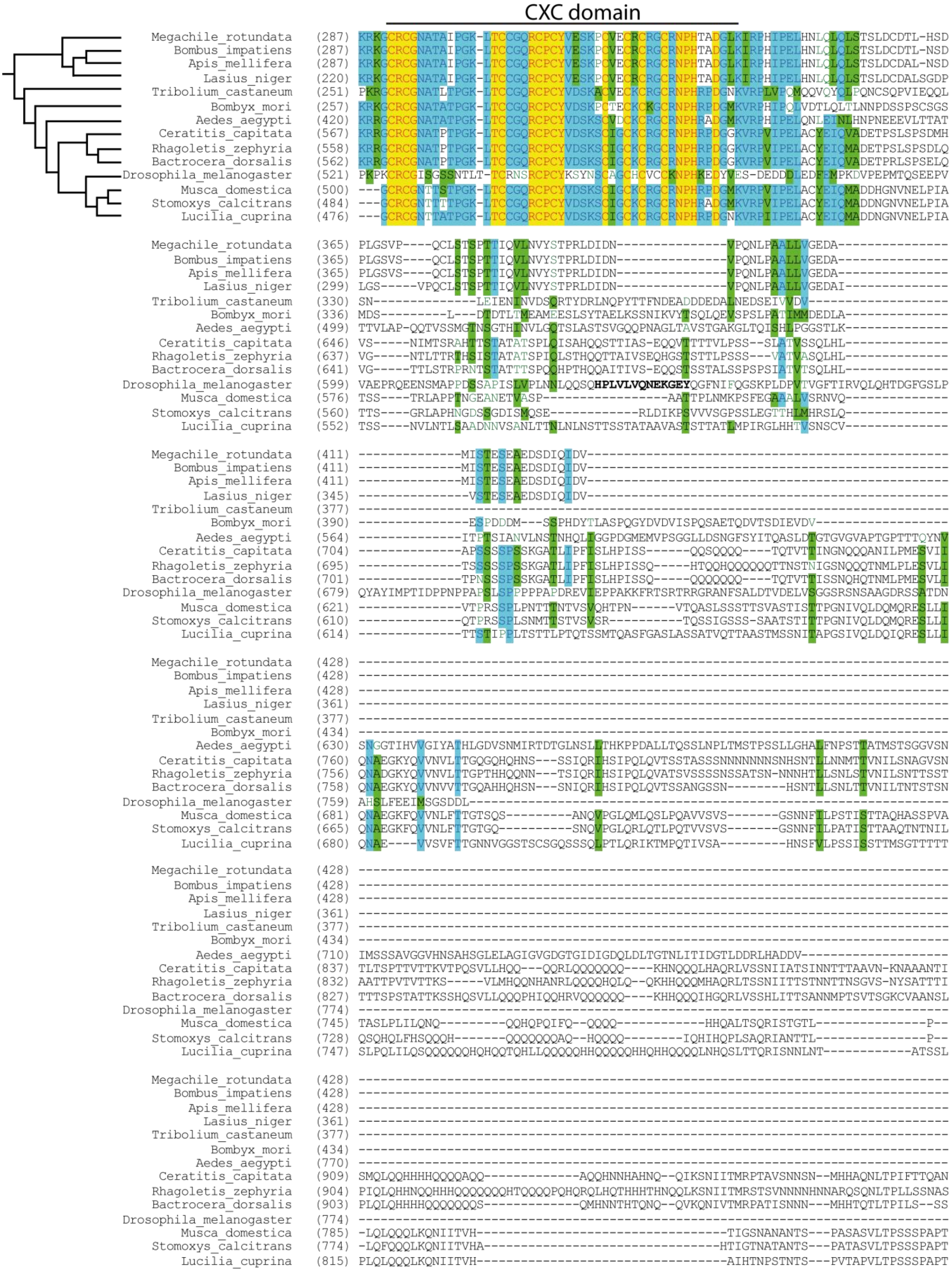

**Supplementary Figure S13. (A)** Sequence alignment of CLAMP N-terminal C2H2 zinc-fingers from honey bee (*Apis mellifera*) and *D. melanogaster*. Typical DNA-binding residues are shown; red asterisks mark the residues of *D. melanogaster* CLAMP displaying the largest chemical shift perturbations upon binding the MSL2 peptide. **(B)** Interaction between GST-tagged CLAMP and 6xHis-tagged MSL2 proteins from honey bee (amCLAMP and amMSL2) and *D. melanogaster* studied with GST/6xHis-pulldown.

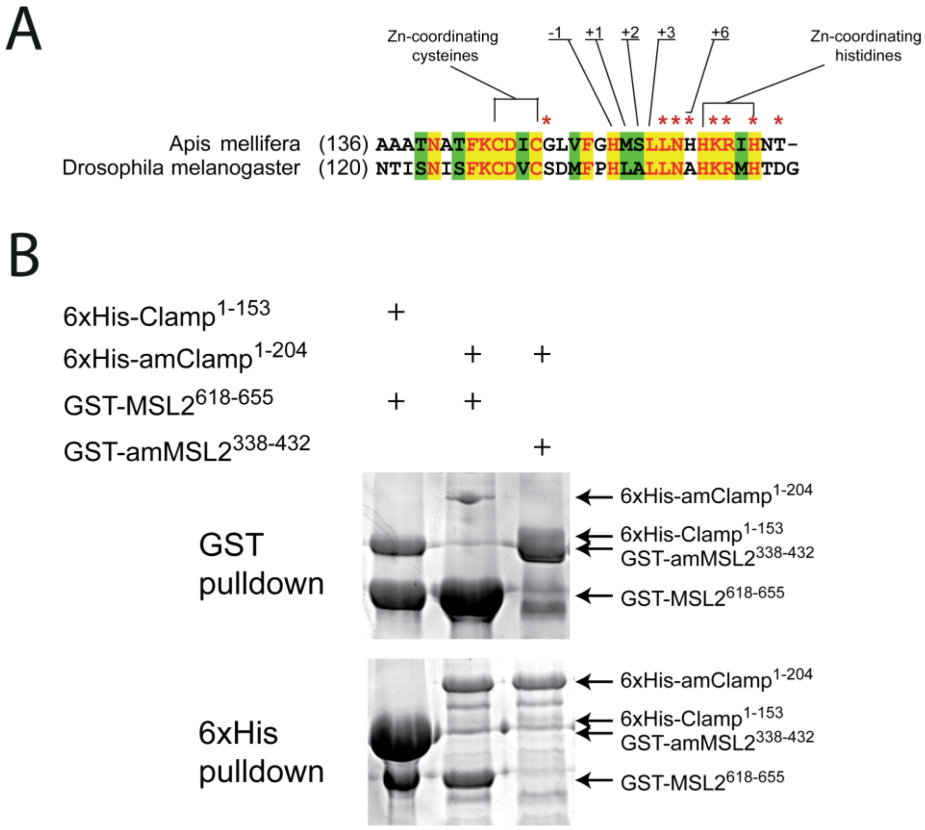

**Supplementary Figure S14**. **(A)** Western blots of protein extracts from transgenic flies expressing wild-type and mutant CLAMP proteins (left) and wild-type and mutant MSL2 proteins (right). **(B)** Effect of single amino-acid substitutions in the FLAG-tagged MSL2 protein on DCC recruitment shown by immunostaining of polytene chromosomes with MSL2 antibodies in females**.** Scale bar is 20 μm.

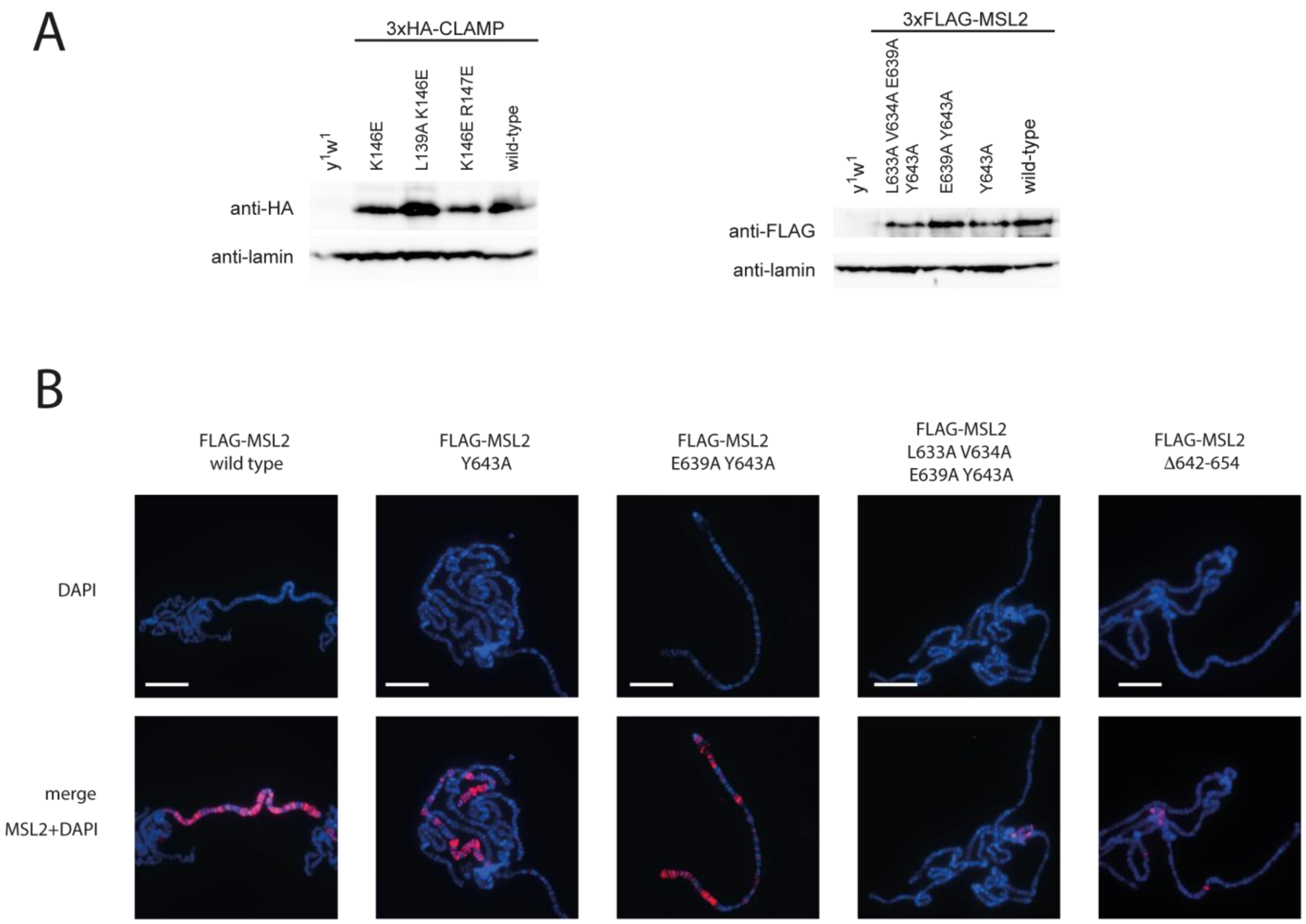

**Supplementary Figure S15.** Schematic representations of interactions mediated by C2H2 zinc-fingers: intramolecular (upper row), DNA interaction and interactions mediated by UBZ-type fingers (lower row). Cartoons were drawn according to following structures (PDB ID): 2GLI (Gli F1 – F2), 1NCS (Swi5), 6DF5 (Kaiso F1 and F3), 1AAY (Zif268-DNA), 3WWQ (FAAP20 UBZ – Ubiquitin), 3VHT (WRNIP UBZ – Ubiquitin).

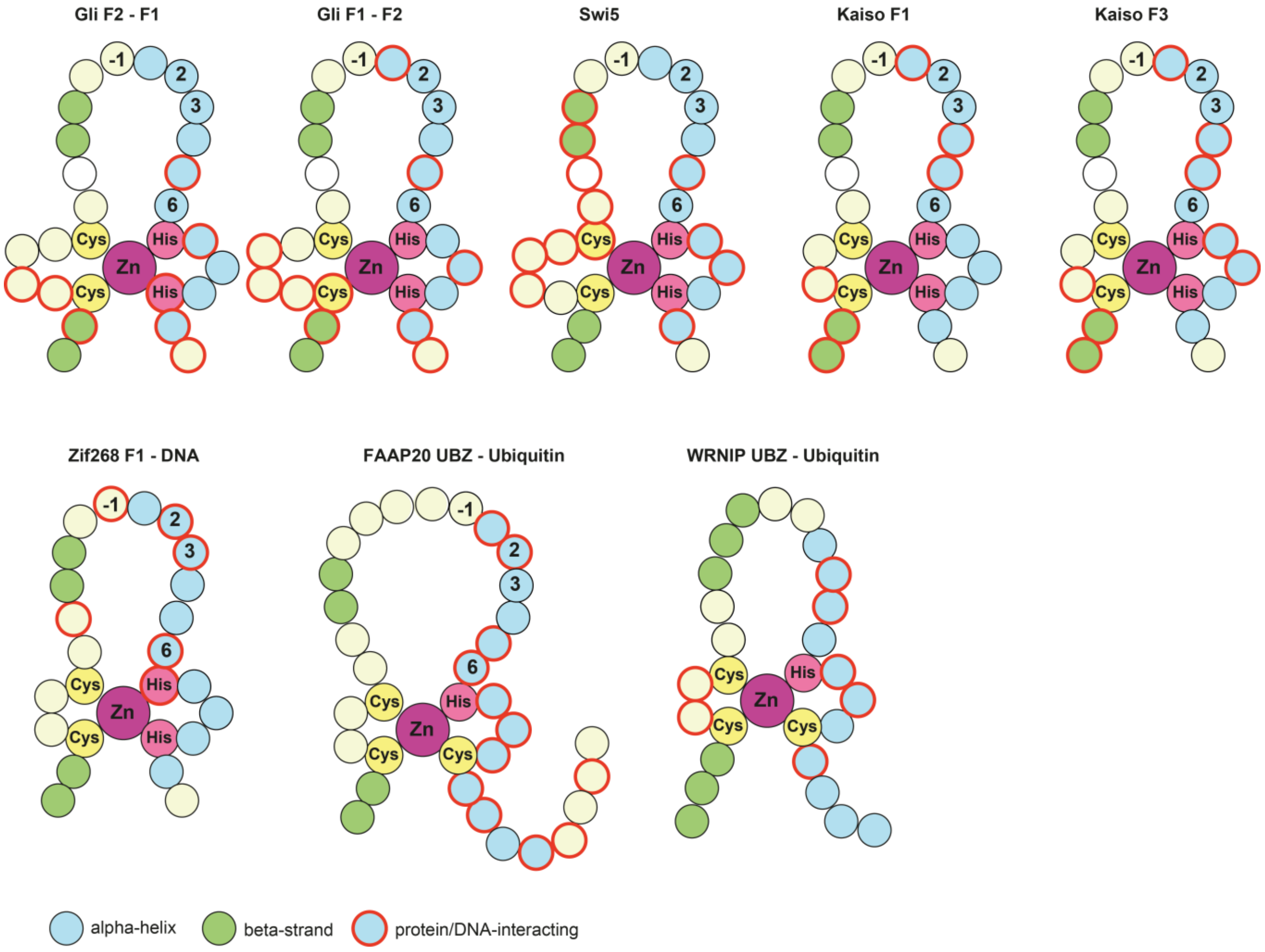

**Supplementary Figure S16.** Ramachandran plot for CLAMP$^{87\text{-}153}$ NMR structure. Glycine residues are represented as white triangles, all other residues are shown as white squares. The general regions corresponding to highly preferred observations are colored in red, moderate and less preferred regions are shown in yellow and ochre, and restricted area is colored in white.

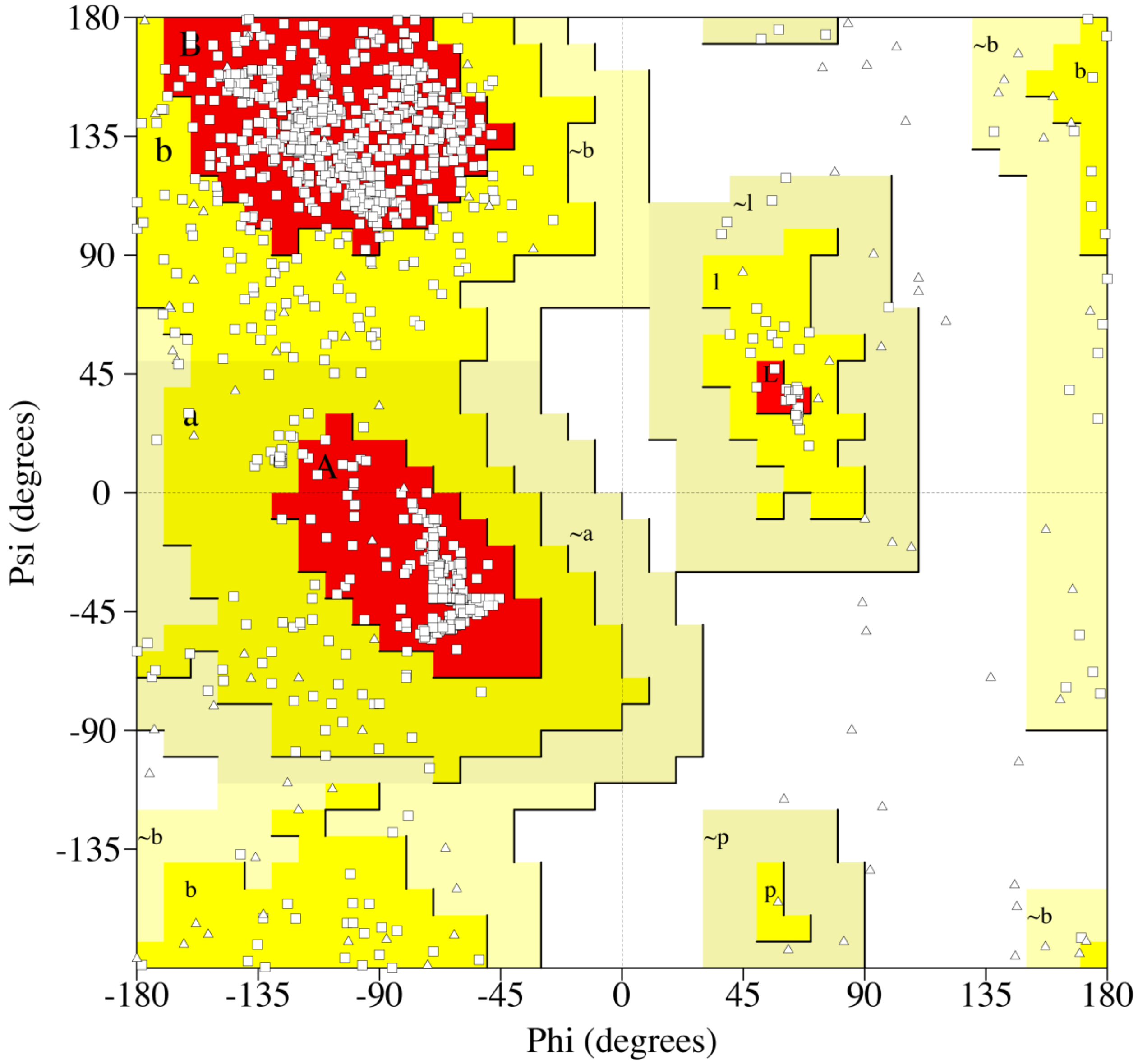

**SUPPLEMENTARY REFERENCES**


1. Bischof, J., Maeda, R.K., Hediger, M., Karch, F. and Basler, K. (2007) An optimized transgenesis system for Drosophila using germ-line-specific phiC31 integrases. *Proc Natl Acad Sci U S A*, **104**, 3312-3317.
2. Sabirov, M., Kyrchanova, O., Pokholkova, G.V., Bonchuk, A., Klimenko, N., Belova, E., Zhimulev, I.F., Maksimenko, O. and Georgiev, P. (2021) Mechanism and functional role of the interaction between CP190 and the architectural protein Pita in Drosophila melanogaster. *Epigenetics Chromatin*, **14**, 16.
3. Tikhonova, E., Fedotova, A., Bonchuk, A., Mogila, V., Larschan, E.N., Georgiev, P. and Maksimenko, O. (2019) The simultaneous interaction of MSL2 with CLAMP and DNA provides redundancy in the initiation of dosage compensation in Drosophila males. *Development*, **146**.
4. El-Gebali, S., Mistry, J., Bateman, A., Eddy, S.R., Luciani, A., Potter, S.C., Qureshi, M., Richardson, L.J., Salazar, G.A., Smart, A. *et al.* (2019) The Pfam protein families database in 2019. *Nucleic Acids Res*, **47**, D427-D432.
5. Vandevenne, M., Jacques, D.A., Artuz, C., Nguyen, C.D., Kwan, A.H., Segal, D.J., Matthews, J.M., Crossley, M., Guss, J.M. and Mackay, J.P. (2013) New insights into DNA recognition by zinc fingers revealed by structural analysis of the oncoprotein ZNF217. *J Biol Chem*, **288**, 10616-10627.
6. Gupta, A., Christensen, R.G., Bell, H.A., Goodwin, M., Patel, R.Y., Pandey, M., Enuameh, M.S., Rayla, A.L., Zhu, C., Thibodeau-Beganny, S. *et al.* (2014) An improved predictive recognition model for Cys(2)-His(2) zinc finger proteins. *Nucleic Acids Res*, **42**, 4800-4812.
7. Persikov, A.V., Wetzel, J.L., Rowland, E.F., Oakes, B.L., Xu, D.J., Singh, M. and Noyes, M.B. (2015) A systematic survey of the Cys2His2 zinc finger DNA-binding landscape. *Nucleic Acids Res*, **43**, 1965-1984.
8. Pavletich, N.P. and Pabo, C.O. (1991) Zinc finger-DNA recognition: crystal structure of a Zif268-DNA complex at 2.1 A. *Science*, **252**, 809-817.
9. Persikov, A.V. and Singh, M. (2014) De novo prediction of DNA-binding specificities for Cys2His2 zinc finger proteins. *Nucleic Acids Res*, **42**, 97-108.
10. Delaglio, F., Grzesiek, S., Vuister, G.W., Zhu, G., Pfeifer, J. and Bax, A. (1995) NMRPipe: a multidimensional spectral processing system based on UNIX pipes. *J Biomol NMR*, **6**, 277-293.
11. Lee, W., Tonelli, M. and Markley, J.L. (2015) NMRFAM-SPARKY: enhanced software for biomolecular NMR spectroscopy. *Bioinformatics*, **31**, 1325-1327.
12. Shen, Y., Delaglio, F., Cornilescu, G. and Bax, A. (2009) TALOS plus : a hybrid method for predicting protein backbone torsion angles from NMR chemical shifts. *Journal of Biomolecular Nmr*, **44**, 213-223.
13. EngineeringToolBox. (2004), *https://www.engineeringtoolbox.com/water-dynamic-kinematic-viscosity-d_596.html*, Vol. https://www.engineeringtoolbox.com/water-dynamic-kinematic-viscosity-d_596.html.
14. Linge, J.P., Habeck, M., Rieping, W. and Nilges, M. (2003) ARIA: automated NOE assignment and NMR structure calculation. *Bioinformatics*, **19**, 315-316.
15. Brunger, A.T., Adams, P.D., Clore, G.M., DeLano, W.L., Gros, P., Grosse-Kunstleve, R.W., Jiang, J.S., Kuszewski, J., Nilges, M., Pannu, N.S. *et al.* (1998) Crystallography & NMR system: A new software suite for macromolecular structure determination. *Acta Crystallogr D*, **54**, 905-921.
16. Shen, Y., Vernon, R., Baker, D. and Bax, A. (2009) De novo protein structure generation from incomplete chemical shift assignments. *Journal of Biomolecular Nmr*, **43**, 63-78.
17. Schanda, P., Kupce, E. and Brutscher, B. (2005) SOFAST-HMQC experiments for recording two-dimensional heteronuclear correlation spectra of proteins within a few seconds. *J Biomol NMR*, **33**, 199-211.
18. Polshakov, V.I., Batuev, E. A., Mantsyzov, A. B. (2019) NMR screening and studies of target–ligand interactions. *Russ Chem Rev.* , **88**, 59-98.
19. Misof, B., Liu, S., Meusemann, K., Peters, R.S., Donath, A., Mayer, C., Frandsen, P.B., Ware, J., Flouri, T., Beutel, R.G. *et al.* (2014) Phylogenomics resolves the timing and pattern of insect evolution. *Science*, **346**, 763-767.

20. Wheeler, T.J., Clements, J. and Finn, R.D. (2014) Skylign: a tool for creating informative, interactive logos representing sequence alignments and profile hidden Markov models. *BMC Bioinformatics*, **15**, 7.
